# Parvalbumin and somatostatin interneurons support distinct computations during auditory scene analysis

**DOI:** 10.64898/2026.07.08.736456

**Authors:** Zhili Qu, Jian Carlo Nocon, Kamal Sen, Howard J. Gritton

## Abstract

Auditory scene analysis requires cortical circuits to preserve target-sound representations while suppressing interference from competing sound sources. Although spatial separation between target and masker sounds can substantially improve perception, the inhibitory circuit mechanisms that transform spatial separation into improved target sound representations remain poorly understood. Here, we used a multi-speaker auditory environment to present target sounds alone (clean) or together with spatially distributed competing white noise (masked) trials while recording auditory cortical activity in awake mice. By combining electrophysiology, cell-type-specific optogenetic suppression, and spike-distance-based neural classifiers, we examined how parvalbumin (PV) and somatostatin (SST) interneurons contribute to spatial processing in complex auditory scenes. PV and SST interneurons made dissociable, context-dependent contributions. PV suppression broadly altered spatial tuning properties, including tuning width, modulation depth, centroid, and sparseness, and reduced target-sound discriminability under clean conditions. In contrast, SST suppression produced comparatively modest effects on spatial tuning but selectively impaired neural discrimination only during masked trials thereby weakening the improvement in target discrimination normally associated with spatial release from masking. Thus, the interneuron population that most strongly altered spatial tuning was not the population that most strongly disrupted masked discrimination. This reveals a dissociation between cortical inhibitory mechanisms supporting spatial receptive-field structure and those supporting target–masker segregation. Together, these findings suggest that PV-mediated inhibition refines the fidelity of auditory cortical representations, whereas SST-mediated inhibition enhances separation of simultaneously spatially competing sound sources, providing complementary inhibitory mechanisms for auditory scene analysis.

## Introduction

A central challenge for auditory perception is maintaining robust representations of behaviorally relevant sounds despite interference from competing acoustic sources. In natural environments, listeners are routinely confronted with complex auditory scenes containing multiple simultaneous sound sources that overlap in time, spectral content, and spatial location. Nevertheless, the auditory system can reliably identify and track target sources while suppressing competing inputs, a remarkable computational ability commonly referred to as auditory scene analysis (Bregman 1990; Willmore et al. 2014). A classic example is the cocktail party problem, where listeners must track the acoustic voice features of a particular speaker in the presence of other competing talkers and background sounds (Cherry 1953). One of the most powerful cues supporting this process is a spatial separation between target and masker sources that aids listeners in detecting and identifying a target more effectively than when the sources are co-located, a phenomenon known as spatial release from masking (Saberi et al. 1991; Bronkhorst 2000; Ihlefeld and Shinn-Cunningham 2008; Kidd et al. 2016). Despite its importance for everyday listening, the neural mechanisms that transform spatial separation into improved representations of attended streams, or targets, remain incompletely understood.

The auditory cortex is thought to play a critical role in solving this problem because it occupies a central position where spatial, spectral, temporal, and behavioral information converge. Previous studies have shown that auditory cortical neurons exhibit strong sensitivity to spatial target–masker configurations and can represent competing sound sources during complex listening tasks (Middlebrooks and Bremen 2013; Maddox et al. 2012; Nocon et al. 2023a). Furthermore, studies in humans have demonstrated that cortical activity selectively tracks attended sound streams and dynamically reorganizes auditory representations according to behavioral demands (Mesgarani and Chang 2012; Zion Golumbic et al. 2013). Together, these findings suggest that the auditory cortex does not merely inherit spatial information from earlier auditory pathways but actively transforms sensory representations to enhance target sounds while suppressing competing sources. However, the circuit mechanisms that support this transformation remain largely unknown.

A growing body of theoretical and experimental work suggests that inhibitory circuits may play a central role in auditory scene analysis. Computational models of spatial target–masker competition predict that inhibitory interactions across location sensitive channels are necessary to generate location-specific enhancements in neural discriminability, referred to as spatial hotspots, during target–masker competition (Dong et al. 2016; Boyd et al. 2025; Nocon et al. 2023a). Similarly, previous physiological studies have shown that inhibitory neurons can profoundly influence sensory tuning, gain control, temporal precision, and signal-to-noise ratio in auditory cortex (Aizenberg et al. 2015; Natan et al. 2015, 2017; Phillips and Hasenstaub 2016). These observations raise the distinct possibility that inhibitory circuits contribute directly to the cortical computations that support spatial release from masking.

In the mammalian cortex, inhibitory neurons comprise approximately 20-30% of the neural population and can be divided into three major non-overlapping classes defined by molecular markers: parvalbumin (PV), somatostatin (SST), and vasoactive intestinal peptide (VIP) neurons (Gonchar and Burkhalter 1997; Yavorska and Wehr 2016). Among the diverse classes of GABAergic interneurons, PV and SST neurons constitute two major, and largely non-overlapping populations, that target excitatory cells but have distinct physiological properties, connectivity patterns, and functional roles (Rudy et al. 2011; Pfeffer et al. 2013). PV neurons primarily target the soma and proximal dendrites of other excitatory neurons and have been implicated in maintaining excitation–inhibition balance, regulating temporal precision, and stabilizing cortical activity (Kubota et al. 2016; Beierlein et al. 2003; Sohal et al. 2009). In auditory cortex, PV neurons sharpen receptive-field structure, enhance temporal coding, and improve sensory encoding reliability (Aizenberg et al. 2015; Jang et al. 2020; Nocon et al. 2023a; Natan et al. 2015; Phillips and Hasenstaub 2016). In contrast, SST neurons primarily target dendrites of pyramidal neurons and have been associated with surround suppression, sustained inhibition, context-dependent sensory processing, and stimulus-specific suppression (Natan et al. 2015, 2017; Lakunina et al. 2020; Muñoz et al. 2017; Chiu et al. 2019; Adesnik et al. 2012; Hendricks et al. 2026). These distinct physiological and anatomical properties suggest that PV and SST neurons may contribute differently to cortical processing during auditory scene analysis.

Despite extensive investigation of PV and SST neurons in shaping frequency tuning, receptive-field structure, gain control, and temporal coding, how these two primary inhibitory populations contribute to auditory spatial processing and target–masker competition remains unresolved. In particular, it is unknown whether the same inhibitory mechanisms that establish spatial receptive-field structure also support the segregation of competing sound sources during auditory scene analysis. To address these questions, we combined a cocktail party-like spatial masking paradigm with electrophysiology, cell-type-specific optogenetic suppression, and spike-distance-based neural classifiers used to decode highly similar target speech envelopes from one another using neuronal spike trains in awake mice. We hypothesized that PV and SST interneurons could putatively support distinct components of auditory scene analysis, with one class contributing to the fidelity of cortical spatial representations while the other contributed to facilitating target–masker segregation during spatial competition. Consistent with this hypothesis, we found that

PV-suppression broadly altered spatial tuning properties while impairing target discrimination predominantly in clean or non-masked conditions, whereas SST suppression produced comparatively modest effects on spatial tuning but impaired target–masker segregation specifically. These results reveal defined roles for PV and SST neurons in supporting auditory spatial tuning and auditory scene analysis.

## Results

To determine how inhibitory circuits contribute to spatial processing during auditory scene analysis, we recorded neuronal activity from primary auditory cortex (A1) while mice were presented with spatially distributed target sounds either alone or in the presence of competing maskers. Because spatial release from masking depends on the ability of cortical circuits to represent both sound location and target identity, we first established the baseline spatial tuning properties of A1 neurons before examining how these representations were altered by cell-type-specific optogenetic suppression. We targeted 32-channel microelectrode arrays to the functionally identified primary auditory cortex (A1) using intrinsic optical imaging (Figure 1a-b) (Narayanan et al. 2023). Tonotopic maps generated from neurovascular/hemodynamic responses to 3kHz, 10kHz, and 20kHz tones were used to identify A1 boundaries prior to implantation (Figure 1a) (Stiebler et al. 1997; Guo et al. 2012).

**Figure 1.**
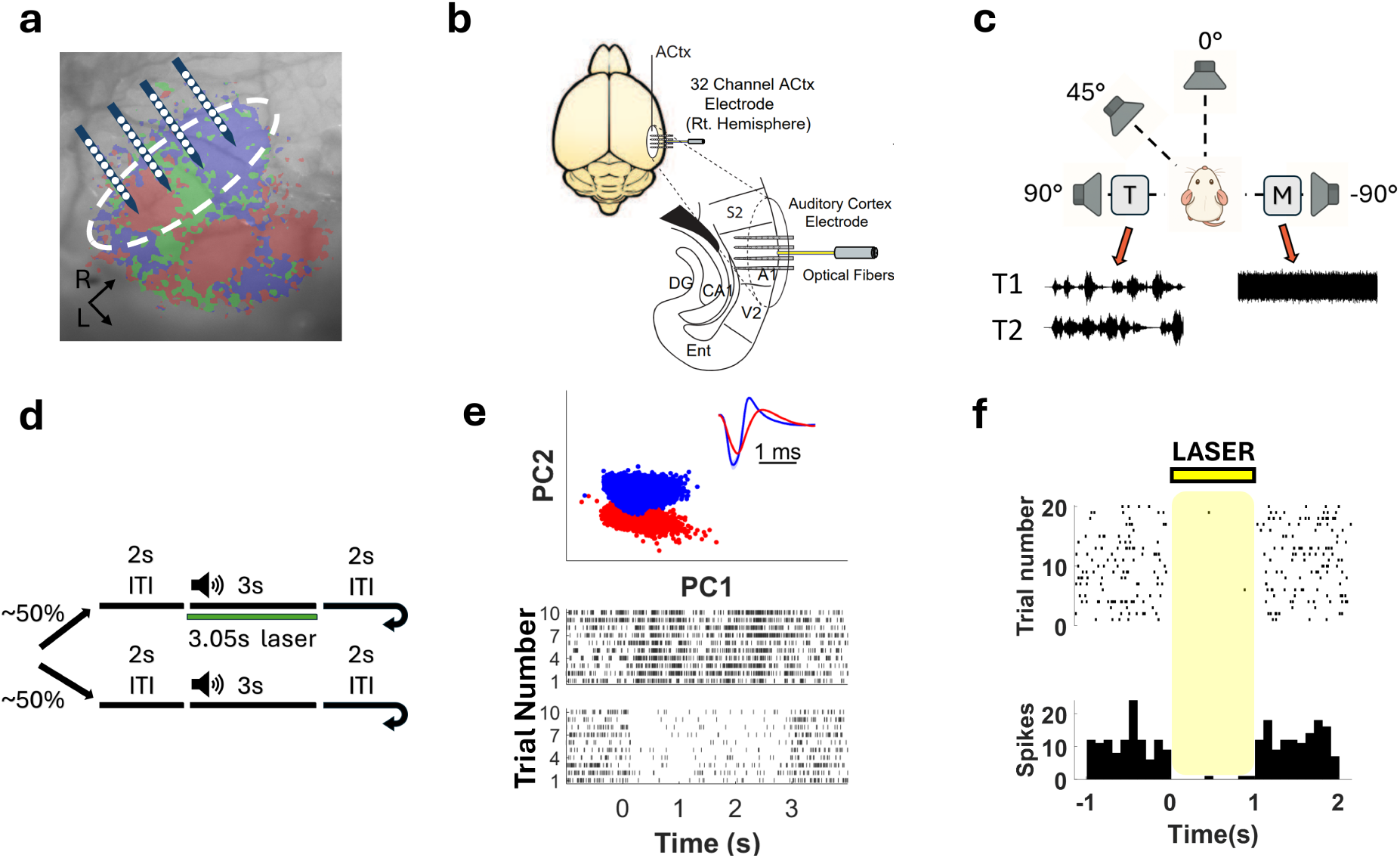
Experimental design and auditory cortical recordings during spatial target–masker competition. **a.** Representative intrinsic signal imaging map used to identify the tonotopic boundaries of pri­mary auditory cortex (A1) before electrode implantation. (red: 3kHz reactive area, green: 10kHz reactive area, blue: 20kHz reactive area). The dashed white line marks the approximate bound­ary of A1. **b.** Schematic illustrating placement of the 4-shank, 32-channel silicon probe and ad­jacent optical fiber within right A1. Electrode arrays were implanted along the tonotopic axis following intrinsic imaging to sample neuronal activity across the primary auditory cortex while enabling optogenetic suppression of local interneurons. Mouse brain illustration is from Pixta (https://www.pixtastock.com/illustration/67155575). **c.** Schematic of the experimental paradigm. Awake, head-fixed mice were passively exposed to two target sounds (T1, T2) presented alone (clean) or together with a competing masker from four loudspeakers positioned at −90°, 0°, 45°, and 90° relative to the animal. Target stimuli consisted of broadband noise modulated by hu­man speech envelopes, whereas masker stimuli consisted of unmodulated broadband noise. Each animal completed 800 randomly interleaved trials spanning all target, masker, and laser condi­tions. **d.** Trial structure design. Each trial consisted of a 2s intertrial interval followed by a 3s auditory stimulus. 50% of the trials consisted of optogenetic trials, during which a 532 nm laser was activated 50 ms before stimulus onset and remained on throughout stimulus presentation. **e.** Representative spike sorting of simultaneously recorded neurons. (*Top*) Example peri-stimulus responses from two single units isolated using Kilosort and Phy, as well as waveform differences captured in 2D principal component space. (*Bottom*) Spike rasters for each unit during the 3 sec­ond target stimulus. **f.** Example electrophysiological validation of optogenetic suppression. Laser illumination reduced firing rate of putative inhibitory neurons in A1, confirming effective sup­pression of Arch-expressing interneurons.

To probe cortical processing during spatial target–masker competition, we implemented a cocktail party-like experimental paradigm in which target sounds and competing maskers could originate from different spatial locations as in our previous studies (Nocon et al. 2023a) (Figure 1c). Four speakers were positioned around the animal at azimuthal locations of −90°, 0°, 45°, and 90° relative to the recorded right auditory cortex. Target stimuli consisted of broadband noise modulated by human speech envelopes (Rothauser et al. 1969) and were represented by two distinct target waveforms (T1 and T2), whereas competing maskers consisted of unmodulated broadband noise. This design allowed us to independently manipulate target identity, target location, masker location, and interneuron activity while quantifying how spatial separation influences cortical representations of target sounds. Mice were presented with target-only (clean) trials or target-plus-masker (masked) trials during simultaneous electrophysiological recording with or without optogenetic suppression of PV or SST neurons (Figure 1d). Control mice, including PV-Cre and SST-Cre mice that were negative for Arch expression were used in control studies. We recorded spiking activity across the 4 shank 32-channel array for which we sorted single units (SUs) from multi-units (MUs) in A1 (Figure 1e). Electrode and opsin expression in PV-Cre and SST-Cre mice was confirmed histologically at the conclusion of the experiment (Figure S1-S2) in addition to electrophysiological confirmation of light-evoked changes in spiking activity of putative interneurons (Figure 1f).

The optogenetic recording session consisted of 400 trials without light and 400 trials with light with silencing occurring on ∼50% of trials, randomly interleaved, throughout the recording sessions although all possible locations and target/masker combinations are equally represented in the laser and non-laser conditions. Each trial consists of a 2 second inter-trial interval and a 3 second stimulus playback with the laser onset 0.05 second ahead of the sound stimulus onset (Figure 1d). Control mice received the exact same trial configuration and light stimulation but did not express the Arch transgene.

Across all recording sessions, spike sorting (Pachitariu et al. 2024; Rossant et al. 2016) identi­fied 194 auditory responsive single units from a total of 427 clusters that exhibited spiking ac­tivity (see Methods). These units included 55 from PV-Arch, 48 units from SST-Arch animals, 52 from PV-Cre control animals and 39 from SST-Cre controls. Individual units formed well-isolated waveform clusters in two-dimensional principal component space and exhibited distinct stimulus-evoked response patterns (Figure 1e), enabling subsequent analysis of spatial tuning and target discriminability at single neuron resolution.

### Auditory cortical neurons exhibit diverse, but contralaterally-dominated spatial tun­ing

Because spatial release from masking depends on the ability of cortical populations to represent sound-source location, we first characterized the baseline spatial tuning properties of A1 neu­rons. Across the population, neurons exhibited diverse spatial response profiles but were gener­ally biased towards sounds originating from the contralateral hemifield relative to the recording site consistent with previous studies (Figure 2). Representative examples included neurons pre­ferring contralateral (90°), intermediate contralateral (45°), central (0°), and ipsilateral (−90°) speaker locations (Figure 2a–p). At the population level, contralateral and 45°-tuned neurons constituted the largest tuning groups (n = 57 and 26 units, respectively), whereas center- and ipsilateral-tuned neurons were less common (n = 23 and 31 units, respectively), consistent with the expected contralateral bias of unilateral auditory cortical recordings.

**Figure 2.**
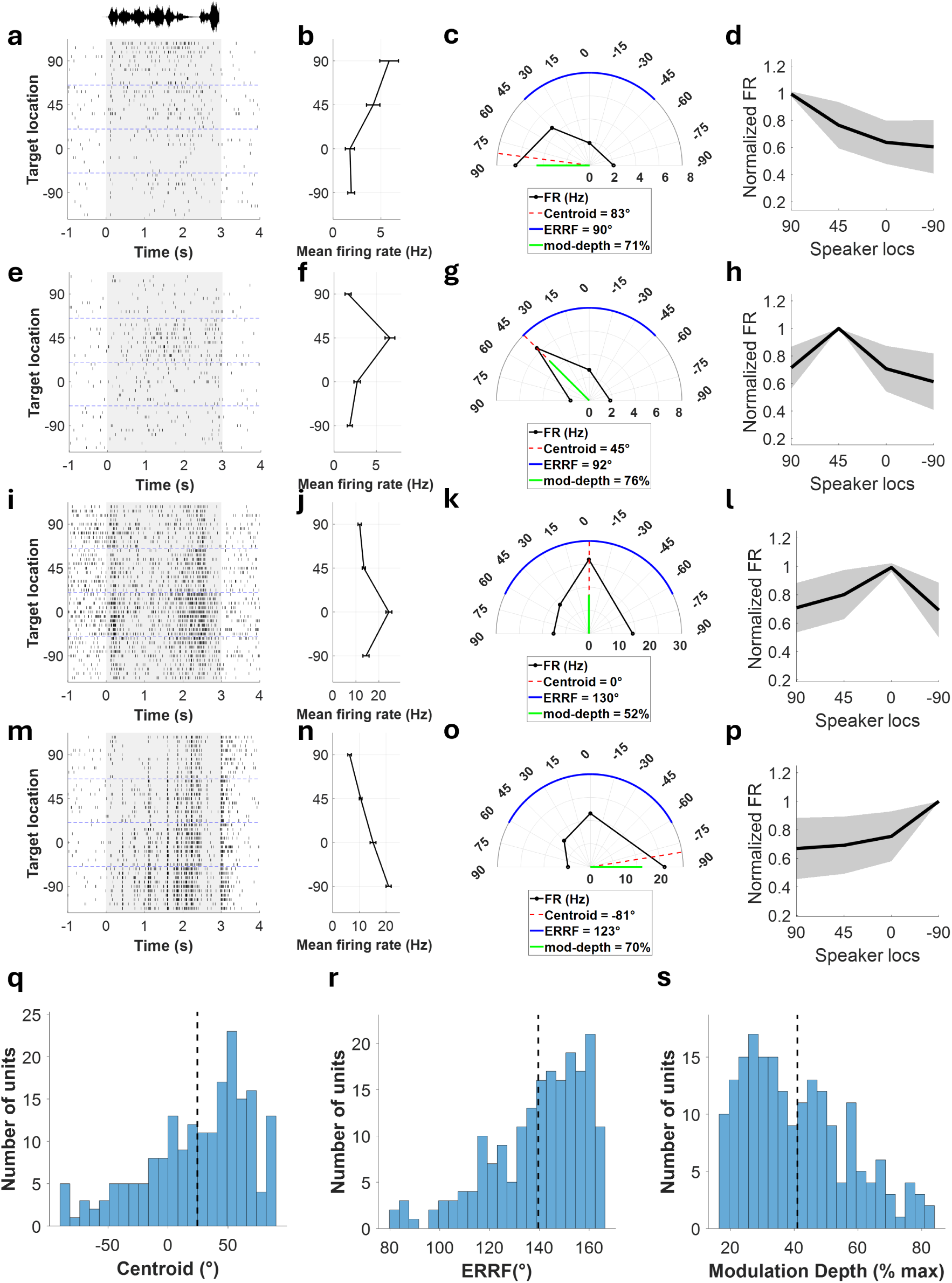
Auditory cortical neurons exhibit diverse but contralaterally biased repre­sentations of auditory space. **a,e,i,m.** Representative peri-stimulus spike raster plots for four example neurons, each preferring one of the four tested speaker locations: contralateral (90°), intermediate contralateral (45°), central (0°), or ipsilateral (−90°). **b,f,j,n.** Average firing-rate tuning curves for the corresponding neurons shown in (a,e,i,m) during auditory stimulus playback from each of the four speaker locations. **c,g,k,o.** Semicircular summaries of spatial tuning prop­erties for the representative neurons shown to the left. Colored overlays indicate the equivalent rectangular receptive field (ERRF: blue), modulation depth (green), and centroid (red). **d,h,l,p.** Population averaged normalized spatial tuning curves for neurons grouped according to preferred sound location (contralateral, 45°, center, or ipsilateral) for all spatially tuned auditory cortical neurons (n = 137) from 27 animals (9 PV-Arch, 8 SST-Arch, 5 PV-Cre and 5 SST-Cre). Gray shading denotes the standard deviation. **q-s.** Histograms of centroids (q), equivalent rectangular receptive field (ERRF) width (r), and modulation depth (s) for all auditory single units. Dashed line represents the mean for each measurement.

To quantify spatial tuning, we isolated single units and quantified each neuron’s preferred sound location (centroid), spatial tuning breadth (equivalent rectangular receptive field: ERRF), and strength of spatial selectivity (modulation depth). (Wood et al. 2019; Lee and Middlebrooks 2011; see Methods). The centroid distribution was skewed toward contralateral locations, with a mean centroid of 24.43° (Figure 2q), consistent with previous reports of opponent spatial coding in auditory cortex (Stecker et al. 2005). Neurons were generally broadly tuned, with an average ERRF width of 139.57°, indicating that many units responded across multiple speaker locations rather than exhibiting narrowly spatial receptive fields (Figure 2r). Nevertheless, substantial spatial selectivity was present in the population, with a mean modulation depth of 40.98% (Figure 2s). Neurons with stronger spatial selectivity typically exhibited higher modulation depth and narrower ERRFs, suggesting a tradeoff between tuning breadth and location specificity across the 180-degree auditory map tested here.

Using a modulation depth criteria of ≥25% (Middlebrooks and Bremen 2013), we classified 137 of 194 auditory-responsive units as spatially tuned. These units spanned all four tuning categories and exhibited substantial diversity in preferred location and tuning breadth. Spatial tuning properties were broadly similar across cortical layers, with no significant differences in modulation depth, centroid, or ERRF between laminar groups (Figure S3). Together, these observations indicate that the mouse auditory cortex contains a heterogeneous but structured representation of auditory space. This representation provides a substrate through which cortical circuits could potentially transform spatial separation into improved target representations during audi­tory scene analysis.

### PV silencing broadly reshapes spatial tuning, whereas SST silencing is selective

Having established that A1 neurons exhibit structured spatial tuning, we next asked whether inhibitory circuits contribute to the maintenance of these spatial representations. To address this question, we optogenetically suppressed either parvalbumin (PV) or somatostatin (SST) interneurons during presentation of clean target sounds and examined how spatial tuning changed at both the single-neuron and population levels. Representative examples illustrate that suppressing either interneuron population increased overall firing rates but produced distinct effects on spatial tuning (Figure 3a-h). In both PV-Arch and SST-Arch animals, optogenetic suppression elevated sound-evoked firing rate activity irrespective of speaker location (Figure 3a-b, e-f). However, the resulting transformations of the spatial tuning curves differed markedly. PV-suppression increased firing rates relatively more uniformly across locations, producing broad upward shifts in tuning curves for this cell (Figure 3c-d) consistent with PV activity contributing to additive gain effects (Lee et al. 2012; Seybold et al. 2015). In contrast, SST suppression preferentially altered firing rate responses near the neuron’s most sensitive location, while largely preserving the overall tuning profile consistent with multiplicative gain effects (El-Boustani and Sur 2014) (Figure 3c-d, g-h). These effects were also evident in the population tuning curves. Across all tuning classes, PV suppression shifted responses upward across speaker locations, whereas SST suppression largely scaled responses while preserving their overall shape (Figure 3i–p). Thus, although both manipulations increased firing rates, they produced qualitatively different effects on the structure of spatial responses.

**Figure 3.**
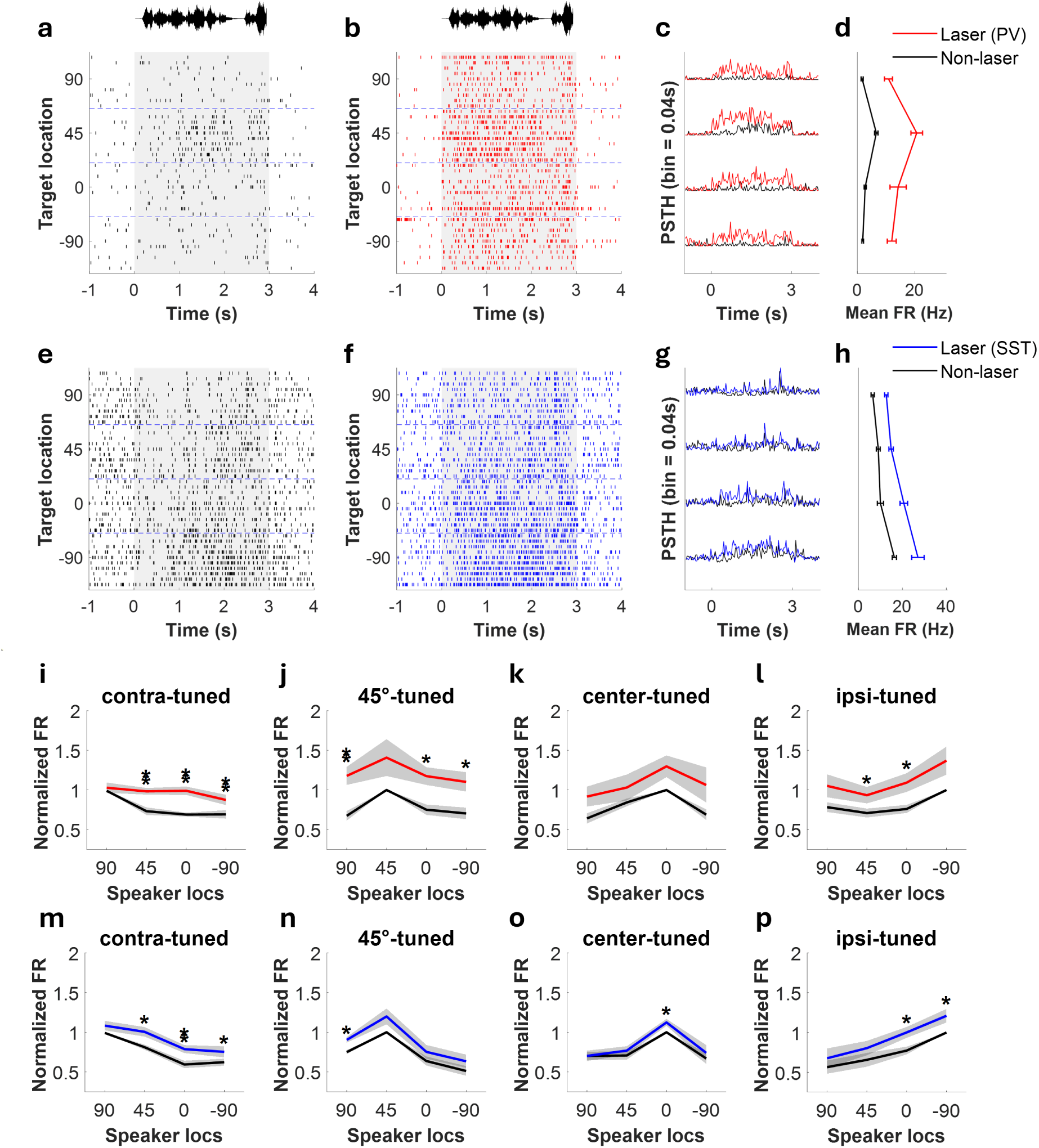
PV and SST interneurons differentially transform auditory spatial representations. **a-b.** Representative auditory cortical neuron recorded from a PV-Arch animal during non-laser (a) and optogenetic laser-suppression trials (b). Peri-stimulus spike rasters show increased firing rate during PV silencing. **c.** Mean firing-rate responses for the neuron shown in (a-b), illustrating increased sound-evoked activity during PV suppression across speaker locations. **d.** Spatial tuning curve for the neuron shown in (a–c), demonstrating a broad upward shift in responses during PV suppression. **e,f.** Representative auditory cortical neuron recorded from an SST-Arch animal during non-laser (e) and optogenetic laser-suppression trials (f). **g.** Mean firing-rate responses for the neuron shown in (e,f), illustrating location-dependent changes in sound-evoked activity during SST suppression. **h.** Spatial tuning curves for the neuron shown in (e–g), demonstrating selective scaling of responses while largely preserving tuning shape. **i-l.** Population average spatial tuning curves from PV-Arch animals (n = 9) grouped according to preferred sound source location (contralateral-tuned n = 15; 45°-tuned n = 9; center-tuned n = 6; ipsi-tuned n = 10). PV suppression produced broad upward shifts in firing rates across spatial tuning classes between non-laser (black) and laser (red) trials. **m-p.** Corresponding population tuning curves from SST-Arch (n = 8) animals (contralateral-tuned n = 16; 45°-tuned n = 5; center-tuned n = 6; ipsi-tuned n = 7). SST suppression produced comparatively selective changes in firing rate while largely preserving the overall structure of the spatial tuning curves between non-laser (black) and laser (blue) trials. Asterisks indicate significant paired comparisons between non-laser and laser conditions (*p < 0.05* ; see Methods).

To quantify these transformations at the population level, we compared firing during non-laser and laser conditions using robust linear regression (RANSAC; Figure S4). PV suppression pri­marily altered the intercept of the non-laser/laser relationship (slope = 1.02, intercept = 0.67), indicating an additive effect on firing rate. In contrast, SST suppression predominantly altered the slope of the relationship (slope = 1.19, intercept = −0.35), indicating a multiplicative trans­formation. These findings suggest that PV and SST interneurons influence auditory spatial repre­sentations through distinct modes of response modulation.

If PV-mediated inhibition contributes broadly to the maintenance of spatial representations, then suppressing PV neurons should alter multiple independent measures of spatial tuning. To test this prediction, we quantified modulation depth, ERRF, and centroid position with and without optogenetic suppression in PV-Arch mice (Figure 4a,e,i) and SST-Arch mice (Figure 4b,f,j). PV suppression altered all three tuning metrics, producing changes in modulation depth (p=1.620 x 10^-6^), ERRF width (p=5.464 x 10^-5^), and centroid (p=0.016: Figure 4c,g,k). In contrast, SST suppression significantly altered ERRF width (p = 0.001) but did not significantly affect modulation depth or centroid position (Figure 4d,h,l). Similar differences were observed when examining response sparseness, or breadth of the responsivity of the cell (Tobin et al. 2025), which was significantly reduced by PV suppression but was largely unaffected by SST suppression (Figure S5). The detailed statistics on how these subpopulations change their spatial tuning can be found in the supplementary information (Figures S6-S8). Importantly, identical laser stimulation in PV-Cre and SST-Cre control animals lacking Arch expression did not systematically alter spatial tuning curves or tuning metrics (Figures S9–S10), indicating that these effects were not attributable to nonspecific optical stimulation.

**Figure 4.**
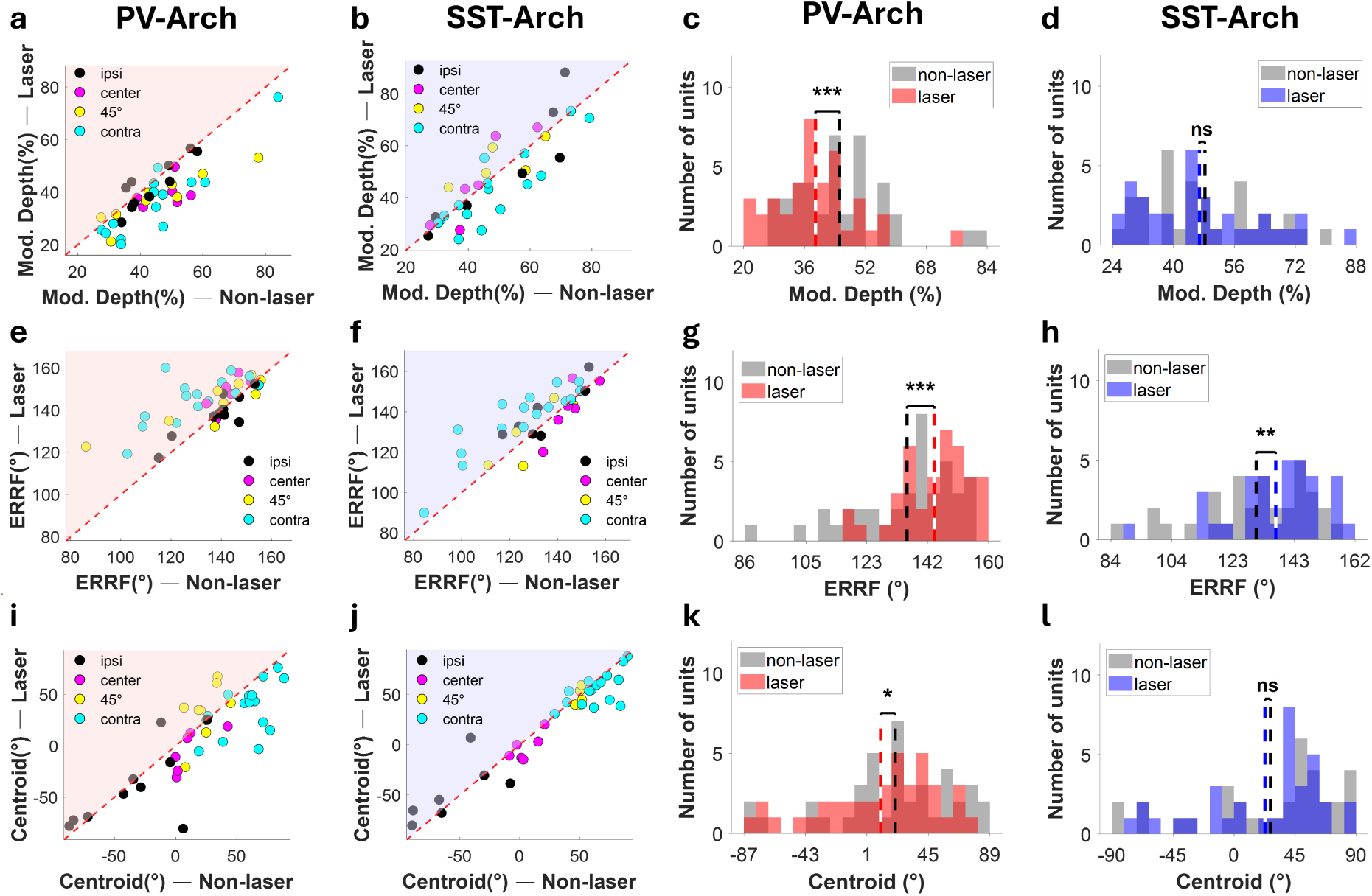
PV suppression broadly alters spatial tuning whereas SST suppression selectively regulates tuning breadth. **a,e,i.** Changes in modulation depth (a), equivalent rectangular receptive field (ERRF; e), and response centroid (i) for individual spatially tuned neurons (n = 40) recorded from PV-Arch animals. Each scatter plot point represents one neuron’s calculated responses from laser and non-laser trials during the recording session. Colors indicate preferred sound location (contralateral, 45°, center, or ipsilateral); Pink shading denotes the region where laser stimulation increases the tuning metric relative to the non-laser condition**. b,f,j.** Corresponding analyses for spatially tuned neurons (n = 34) recorded from SST-Arch animals (cyan shading). **c,g,k.** Population histograms comparing modulation depth (c), ERRF width (g), and centroid position (k) between non-laser (gray) and PV laser-suppression (red) trials. PV suppression significantly altered all three spatial tuning metrics (modulation depth: t(39) = 5.64, p = 1.620 x 10^-6^; ERRF: t(39) = −4.53, p = 5.464 x 10^-5^; and centroid: t(39) = 2.51, p =0.0162). **d,h,l.** Population histogram distributions comparing modulation depth (d), ERRF width (h), and centroid position (l) between non-laser (gray) and SST laser-suppression (blue) trials. SST suppression significantly increased ERRF width (t(33) = −3.59, p = 0.001) but did not significantly alter modulation depth or centroid position. Statistical comparisons were performed using paired *t*-tests (see Methods).

Together, these results demonstrate that PV-mediated inhibition plays a broad role in maintaining the structure of auditory spatial representations, whereas SST-mediated inhibition exerts a more selective influence with impacts targeted to areas of greater spatial tuning sensitivity. Despite producing similar increases in firing rate, suppression of PV neurons altered multiple independent dimensions of spatial tuning, whereas suppression of SST neurons primarily affected tuning breadth. These observations suggest that PV circuits contribute substantially to the organization and stability of cortical spatial representations and raise the possibility that PV and SST interneurons are well-aligned to play dissociable roles in complex acoustic scenes that in­volve competing sound sources.

### Silencing PV or SST interneurons impairs discrimination of target sounds in complex auditory scenes

The preceding results demonstrate that PV suppression produces substantially larger disruptions of spatial tuning than SST suppression. If spatial receptive-field structure directly determines target discriminability, then it is likely that PV suppression would produce the largest deficits in neural discrimination. To explore this, we next asked how suppressing PV or SST interneurons affects the ability of auditory cortical neurons to discriminate between target sounds under clean or masked conditions.

To do this, we used a SPIKE-distance classifier that incorporates both spike timing and firing rate information (Satuvuori and Kreuz 2018; Kreuz et al. 2013). Discrimination performance was measured by comparing spike trains evoked by two highly similar target stimuli (T1 and T2) through a template-matching approach and determining how accurately individual trials could be classified based on their distance from target-specific templates (see Methods). Because SPIKE- distance is sensitive to both temporal precision and response reliability, it provides a useful measure of the quality of cortical representations under different listening conditions. We had previously used this same metric to demonstrate that PV neurons improve neural discrimination of temporally modulated sounds by enhancing temporal coding fidelity and reducing cortical noise (Nocon et al. 2023a). Here, we utilized this same analysis to compare the contributions of PV and SST interneurons during spatial target-masker competition.

For each neuron, discrimination performance was computed across 20 configurations, consisting of four clean target-only conditions and sixteen target–masker conditions. For the example neuron recorded from a representative PV-Arch animal shown in Figure 5a, the non-laser grid shows that discrimination performance varied across spatial configurations, with clean (*top-row*) and masked configurations (*bottom-4x4 grid*) shown in Figure 5c. The corresponding location of rasters shown in Figure 5a (clean) and Figure 5b (masked) is marked by black outlines in Figure 5c. The raster plots and PSTHs corresponding to these selected configurations show that the neuron exhibited temporally structured, target-specific responses during the clean condition (Figure 5a), yielding high discriminability, whereas the presence of a competing masker degraded the target-specific temporal response structure and was associated with reduced discrimination performance (Figure 5b-c). The laser grid shows discrimination performance for the same neuron during optogenetic suppression, with red outlines marking the same selected spatial configurations (Figure 5d). During laser suppression, responses to the two target sounds became less distinct at these configurations, corresponding to reduced discrimination scores relative to the non-laser trials (Figure 5c–d). For the example neuron recorded from an SST-Arch animal, the same organization is shown, with non-laser and laser grids showing discrimination performance across the 20 spatial configurations and raster plots illustrating the underlying clean and masked responses at selected locations (Figure 5g–j). Together, these examples illustrate how spatial configuration, target-locked temporal response structure, and optogenetic suppression jointly shape target discriminability.

**Figure 5.**
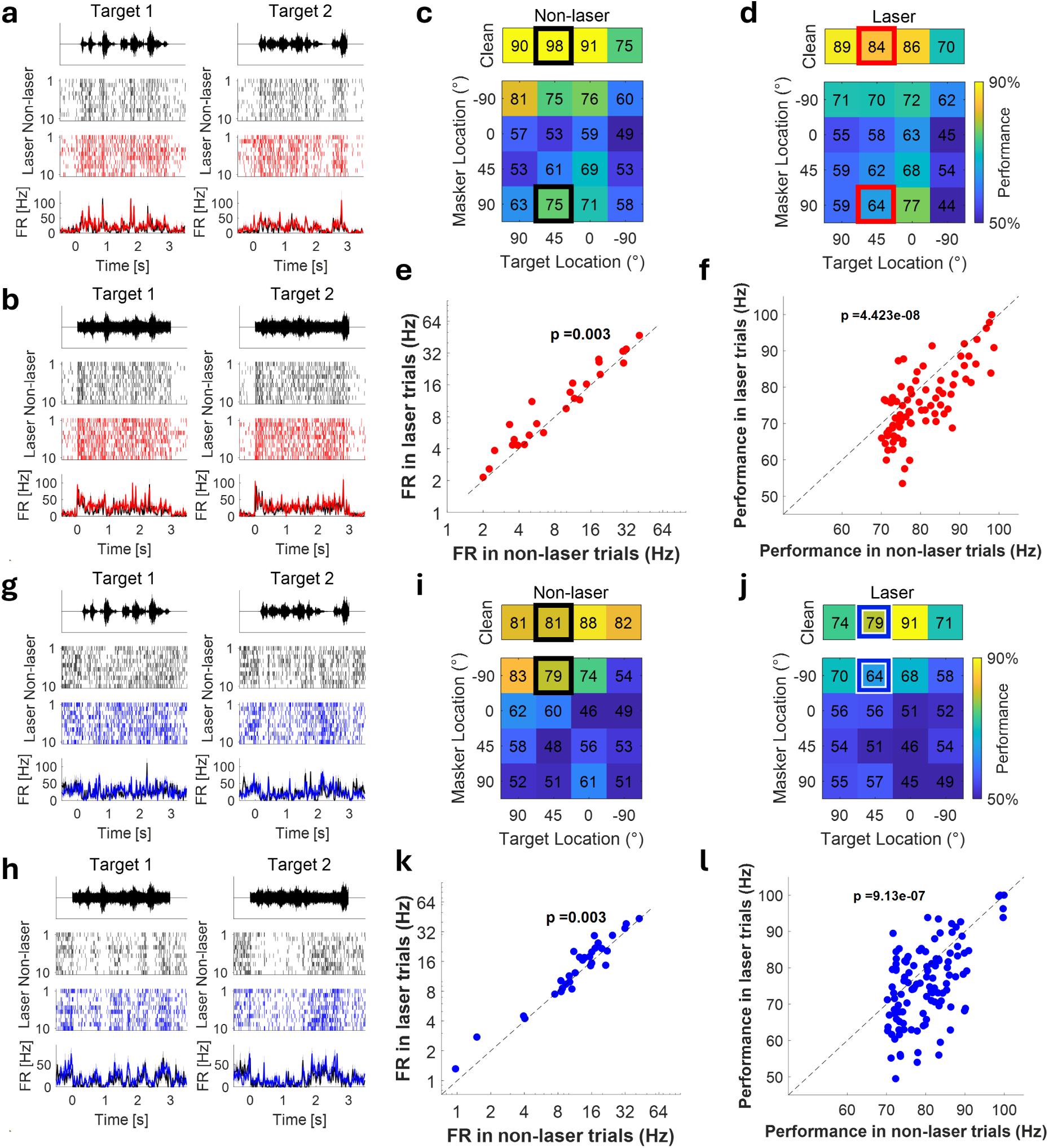
Suppression of PV or SST interneurons impairs neural discrimination at spatial hotspots. **a-b.** Representative responses from an auditory cortical neuron recorded in a PV-Arch animal during clean (a) and masked (b) listening conditions. Raster plots and peri-stimulus time histograms (PSTHs) show responses to the two target sounds (T1 and T2) presented from the clean 45° location from non-laser (top:black) and laser (bottom:red) trials shown in (a). Spiking activity for the combination of target (45°) and masker (90°) sounds is shown in (b). During clean trials, the neuron reliably followed the temporal structure of the target sounds, whereas the addition of a competing masker reduced target-specific response structure**. c-d.** Neural discrimination performance using a template-matching approach (see “Neural discriminability using SPIKE-distance”) for the neuron shown in (a-b) across all target-only and target-masker spatial configurations during non-laser (c) or laser trials (d). Each grid summarizes SPIKE-distance-based discrimination performance for the target alone and target–masker spatial configuration. Spatial configurations with discrimination performance ≥70% and Cohen’s *d* ≥1 were classified as hotspots. Discriminability is generally higher in the top four grids for clean trials, whereas the lower 16 grids represent largely lower performance masked trials. **e.** Scatter plots of average firing rates of hotspot-containing neurons recorded from PV-Arch animals during laser and non-laser trials for auditory neurons that show at least one hotspot (n = 25). PV suppression significantly increased sound-evoked firing rates (t(24) = −3.27, p = 0.003). **f.** Scatter plots of SPIKE-distance-based neural discrimination measured at hotspot configuration with and without PV suppression. Despite increased firing rates, neural discrimination was significantly reduced (n = 76 configurations; t(75) = 6.09, p = 4.423 x 10^-8^). **g-l.** Corresponding analyses for SST-Arch animals. Representative clean (g) and masked (h) responses, spatial discrimination grids during non-laser (i) and SST suppression (j), changes in firing rate (k), and hotspot discrimination performance (l). Similar to PV suppression, SST suppression significantly increased firing rates (n = 32; t(31) = −3.17, p = 0.003) while reducing neural discriminability for hotspots (n = 112 configurations; t(111) = 5.20, p = 9.130 x 10^-7^)

For subsequent population analyses, we defined spatial configurations yielding discrimination per­formance greater than 70% with significant effect sizes as ***hotspots*** (see Methods), representing spatial configurations where cortical neurons could reliably distinguish between the two target sounds. These hotspots provide a framework for assessing how inhibitory circuits contribute to target representation during auditory scene analysis. To determine how inhibitory suppression affects these high-performance hotspot configurations, we first examined changes in firing rate. Suppressing either PV or SST interneurons significantly increased the firing rates of hotspot-containing neurons (PV: *p* = 0.003; SST: *p* = 0.003; Figure 5e,k). However, this increase in ac­tivity associated with suppression of either interneuron population significantly reduced neural discrimination across all hotspot locations (PV: *p* = 4.42 × 10^-8^; SST: *p* = 9.13 × 10^-7^; Figure 5f,l) for the entire population of spatially tuned cells. Thus, disrupting inhibitory circuits increased sound-evoked firing rates while simultaneously degrading neural discrimination, indicating that the quality of cortical representations depends not simply on response magnitude but on the precise spatiotemporal structure of neuronal activity.

Although both manipulations impaired discrimination, the relative contributions of PV and SST circuits under clean and masked listening conditions remained unclear. We therefore next examined whether these interneuron populations differentially support target discrimination across spatial configurations and varying levels of target-masker competition.

### Spatial tuning and target–masker discrimination are differentially affected by PV and SST suppression

Although suppression of both PV and SST interneurons impaired discrimination at hotspot configurations (Figure 5), these analyses did not reveal whether the two interneuron populations contribute similarly across different listening conditions. Given that PV suppression produced substantially larger disruptions of spatial tuning than SST suppression, we next asked whether PV inhibition would also produce the largest deficits in neural discrimination during auditory scene analysis.

To address this question, we calculated average discrimination performance across all auditory neurons exhibiting at least one hotspot in both clean and masked conditions (Figure 6a-f). In PV-Arch animals, the largest decline in discrimination associated with optogenetic suppression was observed during clean trials (Figure 6c). In contrast, SST suppression produced comparatively larger declines in discrimination performance during masked trials, particularly within configurations where the target occupied an equal or more contralateral location relative to the masker (Figure 6f). To quantify these effects, we examined neural discrimination separately for clean and masked conditions. During the clean condition, PV suppression significantly reduced SPIKE distance-based target discrimination across all speaker locations (*p* = 0.003; Figure 6g), indicating that PV-mediated inhibition contributes to maintaining accurate cortical representations of isolated sound sources. However, PV suppression did not significantly affect discrimination in the masked condition (*p* = 0.681; Figure 6h). In contrast, SST suppression did not significantly affect discrimination during the clean condition (*p* = 0.327; Figure 6i), but significantly impaired discrimination during masked conditions (*p*=0.039; Figure 6j). Neither effect was observed in PV-Cre and SST-Cre control animals (Figure S11). Thus, the interneuron population that produced the largest disruptions of spatial tuning did not produce the largest disruption of target discrimination during spatial competition.

**Figure 6.**
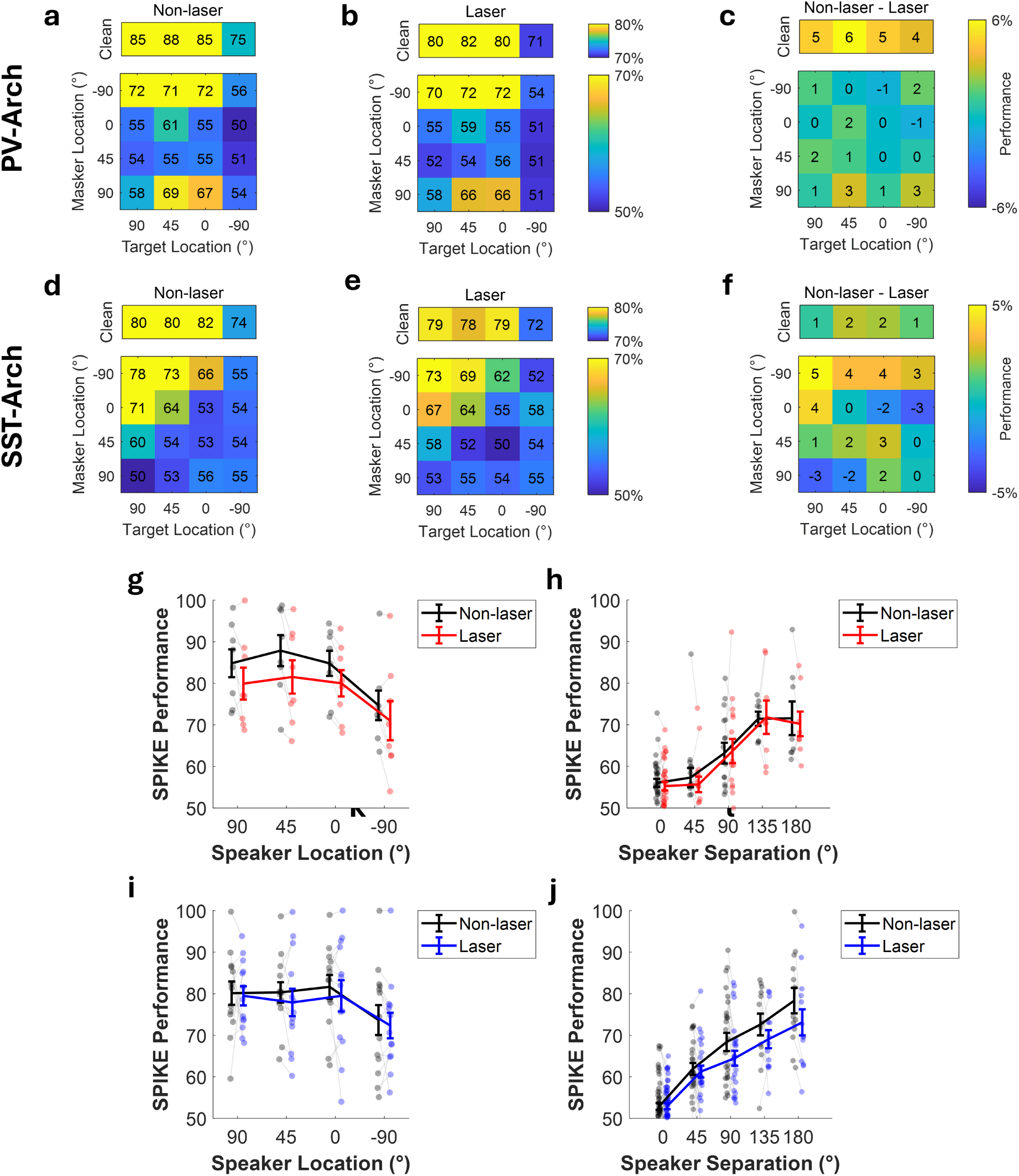
PV and SST interneurons differentially support distinct components of auditory scene analysis. **a-b.** Average SPIKE-distance discrimination performance across all auditory units exhibiting at least one hotspot in both clean and masked conditions from PV-Arch animals during non-laser (a) and PV suppression (b) trials. **c.** Difference map illustrating the effect of PV suppression on the average neural discrimination performance across all target–masker spatial configurations between non-laser (a) and laser (b) trials. PV suppression predominantly reduced discrimination during clean conditions. **d–e.** Corresponding discrimination performance for hotspot-containing neurons from SST-Arch animals during non-laser (d) and laser (e) trials. **f**. Difference map illustrating the average effect of SST suppression on performance across target–masker configurations. SST suppression preferentially reduced discrimination during masked conditions, particularly for configurations in which the target occupied an equal or more contralateral position than the masker. **g.** Scatter plot of average neural discrimination during clean target presentation for PV-Arch animals across four target-only locations. PV suppression significantly reduced SPIKE-distance discrimination (RM-ANOVA: F(1, 7) = 17.80, p = 0.003) across speaker locations between non-laser(black) and laser trials(red). Data plotted as mean ± s.e.m. **h.** Average neural discrimination during masked listening for PV-Arch animals plotted as a function of target-to-masker separation where the target is more contralateral than the masker (non-negative separations). PV suppression did not significantly alter discrimination under masked conditions (Linear mixed-effect model: *β* = 0.0033, SE = 0.0196, t = 0.168, p = 0.867). The x-axis denotes target-to-masker spatial separation. **i.** Average neural discrimination during clean target presentation for SST-Arch animals. SST suppression did not significantly affect clean target discrimination. **j.** Average neural discrimination during masked trials for SST-Arch animals plotted as a function of target-to-masker separation. SST suppression significantly reduced discrimination (RM-ANOVA: F(1, 12) = 5.32, p = 0.039) and weakened the normal improvement associated with increasing target-to-masker separation. The significant laser × separation interaction indicates that SST suppression altered the relationship between spatial separation and neural discrimination rather than producing a uniform reduction in performance (Linear mixed-effects model: *β* = −0.032, SE = 0.014, t = −2.22, p = 0.027).

To further examine the masked condition, we analyzed discrimination as a function of target–masker separation. Following the framework proposed by Maddox et al. (Maddox et al. 2012), target-to-masker separation was defined relative to the recorded hemisphere such that positive values corresponding to configurations in which the target occupied an equal or more contralat­eral position than the masker. Under this population-readout model, increasing target-to-masker separation provides a growing spatial advantage for target encoding within the recorded cortical population. Consistent with previous physiological and behavioral studies (Maddox et al. 2012; Nocon et al. 2023b; Saberi et al. 1991), neural discrimination in non-laser trials improved with increasing target-to-masker separation (Figure 6j). Importantly, SST suppression did not simply reduce discrimination uniformly across masked conditions. Instead, SST suppression preferen­tially weakened the improvement in discrimination that normally accompanied increasing target-to-masker separation. This effect was evident as a reduction in the slope of the discrimination-versus-separation relationship (Figure 6j). A linear mixed-effects model confirmed a significant interaction between laser condition and target-to-masker separation (*p* = 0.027), indicating that SST suppression altered the relationship between spatial separation and neural discriminability rather than producing a constant reduction in performance.

Together, these results reveal a striking dissociation between the effects of inhibitory suppression on spatial tuning and target discrimination. PV suppression produced the larger disruption of spatial receptive-field structure, whereas SST suppression produced the larger deficit in discrim­ination during spatial target–masker competition. These observations suggest that the cortical mechanisms supporting spatial receptive-field organization are dissociable from those support­ing target–masker segregation and indicate that SST-mediated inhibition plays a specialized role during auditory scene analysis when multiple sound sources compete for representation.

## Discussion

In this study, we sought to identify how local inhibitory circuit mechanisms support auditory scene analysis during spatial target–masker competition. By selectively inhibiting parvalbumin (PV) and somatostatin (SST) interneurons using optogenetic suppression during a cocktail-party-like auditory task, we found that each inhibitory population provided distinct contributions to auditory cortical processing. PV suppression broadly altered spatial tuning properties and reduced neural discrimination during clean or target-only conditions, whereas SST suppression produced comparatively modest effects on spatial tuning but selectively impaired neural discrimination during target-masker presentation. This dissociation suggests that the cortical mechanisms responsible for establishing spatial receptive fields are not identical to those that support target–masker segregation during auditory scene analysis. Instead, successful processing of complex auditory scenes appears to depend on multiple inhibitory computations that are recruited in parallel during auditory spatial processing.

Auditory scene analysis requires cortical circuits to preserve information about relevant sounds while suppressing interference from competing sound sources (Bregman 1990; McDermott 2009). Inhibitory interneurons are well positioned to regulate this process as they have been implicated in shaping gain, temporal timing, neural reliability, and enhancing cortical response selectivity (Natan et al. 2015; Nocon et al. 2023a). While previous studies have been essential for elucidating specific roles for PV and SST interneuron populations in shaping frequency tuning, sensory receptive fields, temporal adaptation, gain modulation, surround suppression, and sound detection in noise (Li et al. 2015; Aizenberg et al. 2015; Natan et al. 2015, 2017; Kato et al. 2017; Lakunina et al. 2020, 2022), how each inhibitory subtype contributes to auditory cortical spatial tuning and neural discriminability during spatial target–masker competition remains unresolved. By combining spatially distributed target–masker stimuli with cell-type-specific optogenetic suppression, the present study aimed to dissociate the contributions of each inhibitory cell class during auditory scene analysis.

Our primary observation was a context-dependent division of labor between PV and SST interneurons during auditory spatial processing. Notably, we found that PV-mediated inhibition contributes broadly to the stability and fidelity of cortical spatial representations. Suppressing PV interneurons significantly altered spatial tuning properties across scales in all locations we recorded. This change in spatial sensitivity was observed as alterations in ERRF size, modulation depth, response centroid, and sparseness, indicating that PV circuits influence multiple dimensions of auditory spatial tuning simultaneously. PV neurons have been shown to regulate receptive-field structure, temporal precision, response gain, and sensory encoding reliability in auditory cortex (Aizenberg et al. 2015; Phillips and Hasenstaub 2016; Nocon et al. 2023a). Our results extend these observations by demonstrating that PV-mediated inhibition also contributes substantially to the organization of auditory cortical responses across sound-source locations. Importantly, the effects of PV suppression were not limited to a single tuning metric but were observed across several independent measures of spatial representation and for every location recorded. These observations suggest that PV neurons contribute to maintaining the overall fidelity of cortical representations by preserving the structure and contrast of spatial response patterns.

This broad influence likely arises from the specific manner in which PV circuits modulate cortical responses. PV suppression predominantly shifted the firing-rate intercept producing an additive effect on firing rate across speaker locations. Consistent with previous reports demonstrating context-dependent inhibitory effects (Phillips and Hasenstaub 2016; Natan et al. 2017), this non-selective increase in firing rate elevated responses even at weak or non-preferred locations. Overall, this resulted in impacts on the relative spatial selectivity including alterations in centroid position, sparseness, and modulation depth consistent with previous findings of tuning selectivity (Isaacson and Scanziani 2011). PV suppression also significantly reduced SPIKE-distance-based target discriminability in clean conditions, but this effect was not significant in masked conditions.

In contrast, SST suppression more strongly altered firing-rate slopes. Multiplicative changes can scale responses while largely preserving tuning-curve shape. This additive PV and multiplicative SST pattern differs from the classical frameworks (Wilson et al. 2012; Atallah et al. 2012) and suggests that PV and SST neurons should not be considered as a monolith in support of one type of inhibitory influence on input-output function. Phillips and Hasenstaub (2016) similarly reported that inactivation of SST neurons in the auditory cortex primarily increased response gain, whereas PV inactivation weakened sensory tuning and reduced information transfer, closely matching the response transformations observed here. In addition, SST suppression selectively impaired neural discrimination when target and masker sounds were simultaneously present and spatially separated, while producing comparatively modest effects on clean target discrimination or target-masker configurations with narrow separation. This effect was most pronounced for configurations where the target was collocated with or more contralateral than the masker relative to the recorded hemisphere. In other words, SST activity was most beneficial when the target gained a spatial advantage over the masker, suggesting that SST-mediated inhibition helps convert spatial separation into improved neural discriminability. These results provide a mechanistic explanation of cortical representations of competing sound sources (Maddox et al. 2012; Dong et al. 2016) and reveal a critical role of SST neurons in establishing this representation.

While the PV and SST findings reveal an important role for inhibitory circuits in establishing spatial response structure, the most unexpected observation of this study emerged from the comparison between spatial tuning and neural discrimination for the two cell classes. PV suppression produced substantially larger alterations in classical measures of spatial tuning, including ERRF size, modulation depth, centroid position, and sparseness. However, SST suppression produced the larger deficit in neural discrimination during target–masker competition. If spatial receptive-field structure alone were sufficient to explain target–masker segregation, one might predict that the interneuron population producing the greatest disruption of spatial tuning would also produce the greatest disruption of masked discrimination. Our data did not support this prediction. Instead, they reveal a dissociation between these phenomena, suggesting that the computations supporting auditory scene analysis cannot be fully explained by receptive-field structure alone.

These observations also suggest that successful auditory scene analysis depends in large part on inhibitory mechanisms that are specifically recruited during stimulus competition and are not traditionally captured by conventional measures of spatial tuning. More broadly, these findings indicate that spatial tuning and target–masker segregation represent partially dissociable cortical computations.

This dissociation provides new insight into the functional role of SST interneurons during auditory scene analysis, in particular. Unlike PV suppression, SST suppression had relatively modest effects on spatial tuning but selectively impaired neural discrimination when competing sound sources were simultaneously present. Furthermore, SST suppression weakened the improvement in neural discriminability that normally accompanied increasing target–masker separation. Previous computational models of auditory scene analysis predict that inhibitory interactions operating across spatially sensitive channels would be required to generate location-specific improvements in neural discrimination and the emergence of spatially sensitive hotspots during target–masker competition (Dong et al. 2016; Boyd et al. 2025; Nocon et al. 2023a). By identifying SST neurons as a major contributor to masked-condition discrimination, our findings provide experimental support for this framework while refining its cellular interpretation. Rather than contributing broadly to the establishment of spatial receptive fields, SST-mediated inhibition appears to become particularly important when cortical circuits must resolve competition between multiple simultaneously active sound sources.

The relationship between SST suppression and target–masker separation is especially noteworthy because it connects the present findings to the broader literature on spatial release from masking. Spatial release from masking is one of the most robust perceptual phenomena observed across species, whereby spatial separation between target and masker substantially improves sound detection and identification (Bronkhorst 2015; Ihlefeld and Shinn-Cunningham 2008; Kidd et al. 2016). Although the neural mechanisms underlying this phenomenon remain incompletely understood, our findings suggest that SST-mediated inhibition contributes directly to the cortical circuitry that supports spatial release from masking. Importantly, SST suppression did not simply reduce neural discrimination uniformly across masked conditions. Instead, SST suppression specifically weakened the improvement in neural discrimination that normally accompanies increasing target–masker separation. Thus, SST-mediated inhibition appears to facilitate the transformation of physical spatial separation into a neural advantage for target representation. This interpretation is consistent with the established role of SST neurons in surround suppression, context-dependent sensory filtering, and lateral suppression of competing inputs across cortical systems. (Adesnik et al. 2012; Natan et al. 2015, 2017; Kato et al. 2017; Lakunina et al. 2020). Similar mechanisms have been described in the visual cortex, where SST neurons selectively suppress competing inputs outside the classical receptive field to enhance stimulus selectivity (Adesnik et al. 2012; Karnani et al. 2016; Hendricks et al. 2026). Together, these observations suggest that auditory scene analysis may recruit canonical inhibitory motifs that are shared across sensory modalities.

The present findings also extend our previous work examining the role of PV neurons in complex auditory scenes (Nocon et al. 2023a). In that study, PV suppression degraded neural discrimination through reductions in temporal coding fidelity, spike timing precision, and response reliability. Here, we show that PV and SST neurons contribute to auditory scene analysis through complementary mechanisms. PV neurons support the fidelity of cortical representations by maintaining temporal and spatial response structure, whereas SST neurons contribute more directly to suppressing interference from competing sound sources. Importantly, these findings expand the functional role of PV neurons beyond receptive-field enhancement alone. Together with our previous observations that PV suppression increases cortical noise and disrupts temporal coding (Nocon et al. 2023a), the present results suggest that PV-mediated inhibition refines cortical representations across both temporal and spatial domains. In contrast, SST-mediated inhibition appears to become especially important when cortical circuits must resolve competition between multiple simultaneous sound sources in competition. Viewed together, these findings support a model in which PV and SST interneurons support distinct but complementary components of auditory scene analysis: PV-mediated inhibition preserves representational fidelity, whereas SST-mediated inhibition enables spatial release from masking.

Several limitations should be considered when interpreting these findings. First, although our paradigm captures key acoustic features of the cocktail-party problem, animals were awake while passively exposed to auditory stimuli and the target stimuli lacked explicit behavioral relevance. In natural listening environments, auditory scene analysis is strongly influenced by attention, behavioral goals, and top-down modulation. Previous studies have demonstrated that attentional state can alter spatial sensitivity, enhance auditory discrimination, and promote inhibitory neuron recruitment (Lee and Middlebrooks 2011; Kuchibhotla et al. 2017; Khan et al. 2018; Chou and Sen 2021). Furthermore, functional consequences of interneuron manipulations can also depend strongly on circuit state, stimulus statistics, and experimental context (Lee et al. 2014).

Therefore, future studies combining selective attention tasks with cell-type-specific manipulations will therefore be important for determining how inhibitory circuits contribute to auditory scene analysis during active listening. Second, spatial tuning measurements were based on only four azimuthal locations, providing a relatively coarse estimate of spatial receptive-field structure. More densely sampled speaker arrays may reveal additional effects of inhibitory circuits on spatial representations. Third, our masked-condition analysis used a hemisphere-referenced population readout only from a single hemisphere. Simultaneous bilateral recordings would provide a stronger test of this framework by directly comparing target-favoring and masker-favoring populations contralateral to each recording hemisphere. Finally, because PV and SST neurons operate within highly interconnected cortical networks and converge onto shared pyramidal targets, the effects observed here should be interpreted as circuit-level consequences of sub-type specific perturbations rather than isolated synaptic actions.

In summary, our findings demonstrate that PV and SST interneurons make dissociable, context-dependent contributions to auditory cortical processing during spatial sound processing. More importantly, they reveal that mechanisms supporting spatial receptive-field structure are not those that primarily support target-masker segregation during auditory scene analysis. PV-mediated inhibition plays a prominent role in stabilizing cortical representations across both temporal and spatial domains, whereas SST-mediated inhibition becomes particularly important when competing sound sources must be resolved in complex acoustic environments. By linking distinct interneuron populations to complementary components of target–masker competition, this study advances our understanding of how cortical circuits transform spatial information into robust target representations and provides a circuit-level framework for understanding auditory scene analysis. Difficulties segregating competing sound sources are among the most common complaints reported by hearing-impaired listeners even when detection of isolated sounds remains relatively intact (Shinn-Cunningham and Best 2008; Bronkhorst 2015). Understanding the inhibitory mechanisms that support spatial release from masking may therefore provide biologically grounded principles for enhancing stream segregation in support of cocktail-party-like listening.

## Methods

### Subjects

All procedures involving animals were approved by the Institutional Animal Care and Use Committees (IACUC) at both Boston University and the University of Illinois at Urbana-Champaign. A total of 27 C57BL6/J transgenic mice were used in this study. Original breeding pairs of parvalbumin-Cre (PV-Cre: B6;129P2-Pvalbtm1(cre)Arbr/J), somatostatin-Cre (SST-Cre: STOCK Ssttm2.1(cre)Zjh/J), and Ai40 (Arch: B6.Cg-Gt(ROSA)26Sor tm40.1(CAG-aop3/EGFP)Hze/J) mice were obtained from Jackson Laboratory (Maine), with all breeding conducted in-house. Subjects included both male and female PV-Arch (n = 9) and PV-Cre only (n = 5) mice, as well as SST-Arch (n = 8) and SST-Cre only (n = 5) mice. PV-Arch and SST-Arch mice served as experimental groups, while PV-Cre and SST-Cre mice served as controls. All mice were between 8–12 weeks old on the day of recording.

### Surgery

All mice underwent surgical implantation of a head-plate following our previously published protocol (Nocon et al. 2023a). Briefly, under 2% isoflurane anesthesia, stereotaxic surgery was performed to install a custom head-plate, electrode, and optical fiber. The head-plate was positioned anterior to the bregma and secured to the skull using three stainless steel screws and dental cement. A fourth screw, attached to a metal pin, was placed in the skull above the contralateral cerebellum to serve as a reference. Intrinsic signal imaging was used to delineate the boundary of the primary auditory cortex (A1, right hemisphere) for subsequent electrode placement. A 32-contact linear probe from Neuronexus (model: a 4x8-5mm-100-400-177-CM32), with 100*µ*m spacing between electrode contacts and 400*µ*m spacing between shanks, was precisely inserted into A1 perpendicular to the cortical surface using a stereotaxic arm spanning the low to high frequency axis. Because of the surface curvature of A1, there were minor differences in the absolute depth of each of the four shanks during surgery. As such, the probes were advanced until all electrode contacts were embedded within the cortical tissue (Figure S1) and depth for each shank was recorded. Depth was later confirmed with current source density analysis. Adjacent to the shanks, an optical fiber measuring 200*µ*m in diameter was positioned medially between the two innermost shanks, terminating at the cortical surface (Figure 1b). Following the surgical procedure, mice were allowed a recovery period of 4-7 days before undergoing habituation to being head-fixed as described below.

### Intrinsic signal imaging

Intrinsic signal images were acquired following previously published protocols (Narayanan et al. 2023), using a custom tandem lens macroscope (composed of Nikkor 55 mm 1:2.8 and 85 mm 1:2) and a 16-bit CMOS monochrome camera (BFLY-U3-23S6M-C, FLIR). Prior to the imaging process, chlorprothixene was administered subcutaneously to the mice at a dosage of 1.5 mg/kg body weight, and isoflurane levels were held constant at 1%. The saline wetted skull surface was imaged through a coverslip to increase transparency. The intrinsic signals were captured at a frequency of 20Hz under red light illumination (625nm). The protocol for each trial was composed of a baseline period of 1.5s, a 1s pure tone stimulus at 75 dB SPL across frequencies of 3, 10, or 20kHz, and a 28.5s gap between trials. All stimuli were generated by Tucker Davis Technologies (TDT; Alachua, FL) RZ6 system and played by TDT Multi-Field Magnetic Speakers (MF1) in closed-field operation. Images captured during the response period (0.5s to 2.5s from the sound onset) were averaged and divided by the average pre-stimulus image during the baseline to produce the resultant image. Subsequently, this image was binarized and color-coded to generate the final tonotopy gradient map using customized MATLAB code. A1 regions were then identified manually by their characteristic tonotopic organization.

### Habituation

After the surgical procedures and a recovery period of 4-7 days, mice underwent a handling phase that lasted several days. They were gradually exposed to longer periods of head-fixing to acclimate them to restraint. Each mouse underwent at least six habituation sessions before advancing to the first recording session. To minimize movements while head-fixed, the mice were gently swaddled using lab tissue (Kimwipes: Kimberly-Clark, Irving, TX), secured on both sides with tape. After the head-fixation habituation period, mice were further acclimated to the recording paradigm, during which they were introduced to spatial stimuli, as will be described in the following sections.

### Recording sessions and data acquisition

Recordings were conducted using the TDT RZ10X system within a soundproof chamber (Industrial Acoustics Company, Inc.). Broadband neural signals for all channels were recorded at a sampling rate of 24414.0625 Hz. Local field potentials (LFPs) were digitized at 3051.8 Hz and notch filtered at 60 Hz, before being utilized for current source density analysis. The recording sessions included a mix of optogenetic and non-optogenetic trials, presented in a randomized sequence. Each trial consists of 3 seconds of stimulus playback and 2 seconds of inter-trial interval before the next trial. The mice encountered both target-sound-only (clean) trials and trials with both the target sound and the masker sound co-presented (masked), across various sound source locations. All trials occurred with and without optogenetic silencing in random order. With each trial type being repeated 10 times, the 800 trial session lasted approximately 60 minutes.

### Auditory stimuli

Auditory stimuli were generated in Matlab and delivered through four electrostatic loudspeakers (TDT ES-1; Tucker-Davis Technologies). Target stimuli consisted of broadband white noise amplitude-modulated by temporal envelopes derived from sentences in the Harvard IEEE speech corpus (Rothauser et al. 1969), a source previously used to study auditory scene analysis and the cocktail party problem (Nocon et al. 2023a). Two distinct target stimuli, denoted as T1 and T2, were generated using different speech envelopes while maintaining identical overall stimulus statistics. Competing masker stimuli were unmodulated broadband white noises.

All stimuli were produced with uniform root mean square (RMS) amplitudes and a sampling rate of 195312 Hz to preserve the full frequency range audible to mice. Stimuli were uploaded to an RPvdsEx circuit running on an RZ6 Multi I/O processor (Tucker-Davis Technologies) interfaced with two PM2R multiplexers, allowing independent control of target and masker presentation from each loudspeaker.

During recording sessions, four speakers were positioned 18 cm away from the animal at azimuthal locations of −90°, 0°, 45°, and 90° relative to the right auditory cortex. Speaker outputs were calibrated before each experiment using a conditioning amplifier (Brüel & Kjær, Nærum, Denmark; amplifier model: 2690) and measurement microphone (Brüel & Kjær, Nærum, Denmark; 4939-A-011) to ensure a sound pressure level of 75dB SPL at the animal’s head position. Stimuli were presented for 3s with a 1ms cosine onset and offset ramps.

### Optogenetic stimulation

Optogenetic suppression was achieved using a 532nm DPSS laser (Shanghai Laser Ltd., Shanghai, China; model: GL532T3-200FC) coupled to a 200*µ*m multimode optical fiber (Thorlabs, Newton, NJ; model: M123L02) positioned immediately above the cortical surface and adjacent to the recording array. Laser power was adjusted to 10mW at the tip of the fiber, measured with a calibrated optical power meter (PM100D, Thorlabs, Newton, NJ).

Laser stimulation was synchronized to auditory stimulus presentation through transistor–transistor logic (TTL) pulses generated by the RZ10X recording system. During optogenetic trials, continuous laser illumination began 50ms before sound onset and remained on during the 3s auditory stimulus (Figure 1d), yielding a total illumination duration of 3.05s. Optogenetic and non-optogenetic trials were randomly interleaved throughout the recording session so that every target and masker combination for each target identity was sampled under laser-on and laser-off conditions.

### Histology

Following completion of the experiments, mice were transcardially perfused with 0.01M phosphate buffered saline (PBS) followed by 4% paraformaldehyde (PFA). Brains were postfixed overnight in 4% PFA, in 30% sucrose solution for 24 h, and sectioned coronally (40-50*µ*m) using a freezing microtome. Sections containing the auditory cortex were either Nissl stained to verify electrode placement or processed for immunohistochemistry.

For SST-Arch mice, free-floating sections were permeabilized and blocked in PBS containing 2% normal goat serum and 0.5% Triton X-100 before overnight incubation at 4°C with rabbit anti-SST antibody (Invitrogen, cat. #PA5-82678, 1:1000). Sections were subsequently incubated with goat anti-rabbit Alexa Fluor secondary antibody (Invitrogen, cat. #A-11011, 1:500). Image-iT™ FX Signal Enhancer (Invitrogen, cat. #I36933) was applied according to the manufacturer’s instructions prior to mounting.

For PV-Arch mice, sections were processed using a guinea pig anti-PV primary antibody (SWANT, GP72, 1:1000), followed by goat anti-guinea pig Alexa Fluor 568 secondary antibody (Thermo Fisher Scientific, cat. #A-11075, 1:500). Sections were counterstained with Hoechst 33342 (Thermo Fisher Scientific, cat. #62249, 1:10,000 in 0.01 M PBS) prior to mounting.

All sections were mounted on Fisherbrand Superfrost Plus microscope slides (Fisher Scientific, cat. #12-5550-15). SST-stained tissue was coverslipped using Fluoromount-G with DAPI (Invitrogen, cat. #00-4959-52), whereas PV-stained sections were mounted using ProLong Diamond Antifade Mountant (Thermo Fisher Scientific, cat. #P36965).

### Imaging and Quantification

Arch expression in PV-Arch animals was characterized previously (Nocon et al. 2023a). For SST-Arch mice, fluorescence images were acquired using a Keyence BZ-X800 microscope with a 20× objective. For each section, images were captured at a resolution of 1920 × 1440 pixels, focusing on the plane with the highest visible neuron density. Regions were selected to maintain comparable densities of Arch-GFP–positive cells across animals. To quantify targeting specificity, Arch-GFP expression was assessed within a 400 × 400 *µ*m grid and compared with SST+ immunolabeling . Two regions from two auditory cortical slices were analyzed per animal. Electrode placements were verified independently from Nissl-stained sections imaged with a 2× objective and referenced to a digital stereotaxic atlas.

To standardize fluorescence quantification, we developed a custom MATLAB-based graphical user interface (GUI), **SliceQuant** (Figure S12). SliceQuant provides a semi-automated workflow for measuring fluorescence expression by combining user-defined region-of-interest (ROI) selection with automated cell detection and manual verification. The software identifies Arch-GFP–positive neurons, determines their colocalization with SST-immunopositive cells, and records cell identities, ROI coordinates, and analysis parameters for each image. Adjustable image-processing parameters, including channel-specific contrast, brightness, and size-based object filtering, were used to improve detection while allowing user verification of all identified cells. Quantification results were exported as structured .xlsx files for downstream analysis and reproducibility.

### Spike extraction and clustering

Spike sorting was performed offline using Kilosort2.5 (https://github.com/MouseLand/Kilosort) (Pachitariu et al. 2024), followed by manual curation in Phy (https://github.com/cortex-lab/phy) (Rossant et al. 2016). During curation, clusters representing noise, laser-induced artifacts, or signals common to multiple recording channels were removed. Clusters with overlapping principal component distributions and cross-correlograms indicating a shared refractory period were merged. Curated spike-clusters were subsequently imported into Matlab using the *spikes* toolbox (https://github.com/cortex-lab/spikes) and assigned to recording channels according to the electrode contact with the largest mean spike amplitude. To minimize potential contamination from optical artifacts, spike waveforms exceeding an absolute threshold of 1500 *µ*V or a positive peak threshold of 750 *µ*V were excluded from further analysis. Cluster quality was evaluated using the *sortingQuality* toolbox (https://github.com/cortex-lab/sortingQuality) (Schmitzer-Torbert et al. 2005). Single-units (SUs) were identified using standard quality metrics, including fewer than

5% of inter-spike intervals shorter than 2ms, an isolation distance > 15, and an L-ratio < 0.25. When isolation distance or L-ratio could not be reliably estimated, inter-spike interval violations served as the primary quality criterion, consistent with previous studies (Kvitsiani et al. 2013; Jung et al. 2022). Clusters failing to meet these criteria were classified as multi-units (MUs).

For subsequent analyses, single units were categorized as narrow-spiking when the trough-to-peak waveform duration was <0.5ms, consistent with previous electrophysiology studies identifying excitatory and inhibitory units in mouse auditory cortex (Li et al. 2015). Finally, units were classified as auditory responsive if either their mean or peak firing rate during sound presentation differed significantly from baseline activity (t-test, p-value below 0.05).

### Neural discrimination using SPIKE-distance

Neural discrimination performance was quantified using a template-matching classifier based on the SPIKE-distance metric, following our previous work (Nocon et al. 2023a). SPIKE-distance measures dissimilarity between spike trains by incorporating differences in both spike timing and instantaneous firing rate without requiring user-defined parameters (Satuvuori and Kreuz 2018; Kreuz et al. 2013). Because the metric is sensitive to both temporal precision and response reliability, it provides a robust measure of the quality of cortical representations during auditory scene analysis.

For each spatial target–masker configuration, neural discrimination between the two target stimuli (T1 and T2) was evaluated using template matching. In each of 100 iterations, one of the ten spike trains recorded for each target served as the template, while the remaining 18 spike train trials were compared to the templates to determine the target identity. All possible template pairs were sampled across the 100 iterations, and average classification accuracy was used as the neural discrimination score for that spatial configuration.

The SPIKE-distance between two spike trains was computed as described previously (Satuvuori and Kreuz 2018). Briefly, the instantaneous spike timing difference at time *t* is defined as:

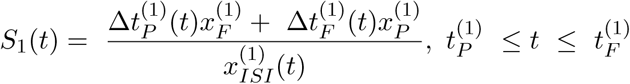

where Δ*t_P_*, Δ*t_F_* denote the distances to the nearest spike in the second train from the preceding and following spikes in the first train, respectively, with *x_F_* and *x_P_* representing the absolute distances from *t* to these spikes, and 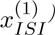 is the interspike interval in the first train. The perspective from the second train, *S_2_(t)*, is calculated by substituting all spike times and intervals with those from the second train. The instantaneous difference between the two trains is then computed as:

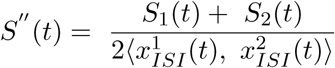

This difference is locally weighted by instantaneous spike rates to account for firing rate differ­ences:

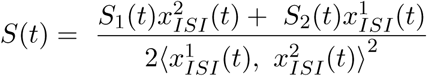

The overall distance between a pair of spike trains is the integral of this dissimilarity profile over time T, where identical spike trains yield a minimum value of 0:

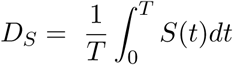

The cSPIKE toolbox (Satuvuori and Kreuz 2018) was used to compute SPIKE-distances for all spike-train pairs across every spatial grid configuration. To identify spatial configurations in which individual neurons reliably discriminated between the two target stimuli, we defined **hotspots** as configurations exhibiting consistently high neural discrimination under non-laser conditions. A hotspot was required to satisfy three criteria: (1) mean discrimination performance during non-laser trials greater than 70%; (2) discrimination significantly greater than chance (*p* < 0.05), determined by comparison with a null distribution obtained by classifying spike trains within the same target stimulus; and (3) a Cohen’s *d* effect size greater than 1 between the observed and null discrimination distributions:

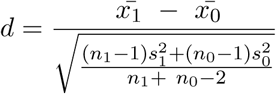

where values with subscript 0 represent the mean, standard deviation, and number of template-matching iterations for the null discrimination distributions, respectively. Spatial configurations in which three or more trials from either target contained no spikes were excluded to avoid unstable estimates of neural discrimination.

### Current source density and Laminae Analysis

Current source density (CSD) analysis was performed to assign recording sites to cortical layers based on the laminar distribution of sound-evoked synaptic currents (Nicholson and Freeman 1975). CSDs were calculated from local field potentials (LFPs) recorded on each shank of the silicon probe. For each recording channel, LFPs were averaged across all trials prior to CSD calculation. The resulting LFP profiles were spatially smoothed across the eight channels of each shank as:

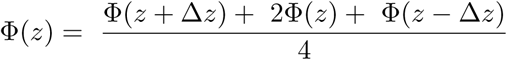

where *z* denotes the depth perpendicular to the cortical surface, Δ*z* is the electrode spacing, and Φis the extracellular potential. CSD was then estimated as:

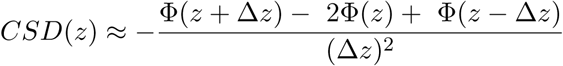

The granular layer (layer 4) was identified as the recording site exhibiting the earliest stimulus-evoked current sink on each shank. When multiple channels exhibited identical sink onset times, layer assignment was based on the earliest sink observed across neighboring recording sites. Recording sites located superficial and deep to layer 4 were classified as layers 2/3 and 5/6, respectively, and were used for the laminar analyses presented in Figure S3.

### Spatial tuning analysis

Spatial tuning curves were constructed by averaging sound-evoked firing rates across repeated presentations of the target stimulus at each of the four loudspeaker locations. To characterize the spatial receptive-field properties of individual neurons, three complementary metrics were calculated: modulation depth, response centroid, and equivalent rectangular receptive field (ERRF), following established methods for auditory spatial coding (Lee and Middlebrooks 2011; Wood et al. 2019).

Spatial selectivity was first quantified using modulation depth, defined as:

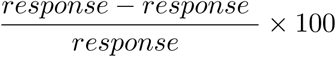

where response (max) and response (min) represent the maximum and minimum firing rates across speaker locations, respectively. Following Middlebrooks and Bremen (2013) but using a reduced threshold appropriate for our four-speaker spatial sampling, neurons with modulation depth ≥ 25% were classified as spatially tuned.

Preferred sound location was quantified using the response centroid. The centroid was calculated by first identifying speaker locations evoking responses greater than or equal to 75% of the unit’s maximum firing rate. These responses were represented as vectors weighted by their corresponding firing rates and the direction of the resultant vector sum was taken as the neuron’s preferred spatial tuning direction.

Spatial tuning breadth was quantified using the equivalent rectangular receptive field (ERRF), defined as the width of a rectangle with an area equal to that under the spatial tuning curve and a height equal to the peak firing rate.

To obtain robust parameter estimates modulation depth, centroid, and ERRF width were estimated using nonparametric bootstrap resampling. Spike counts at each speaker location were resampled with replacement 500 times while preserving the number of trials per location. Each spatial tuning metric was calculated for every bootstrap iteration, and the mean of the resulting distribution was used as the final estimate. The 2.5th and 97.5th percentiles of the bootstrap distributions were taken as the corresponding 95% confidence intervals.

### Response sparseness

As an additional measure of spatial representation, we quantified the sparseness of each neuron’s responses across speaker locations. Whereas ERRF measures the breadth of the spatial receptive field, sparseness quantifies the extent to which a neuron responds selectively to a limited subset of spatial locations. Response sparseness was calculated according to Tobin et al. (2025), defined as:

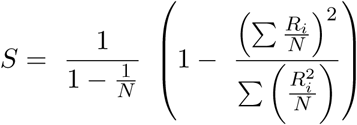

where *R_i_* is the mean firing rate evoked by target sounds presented from the *i-*th speaker location, and *N* is the number of speaker locations. Sparseness values range from 0 to 1, with values approaching 0 indicating broadly distributed responses across speaker locations and values approaching 1 indicating highly selective responses confined to a small number of locations.

### Analysis of multiplicative versus additive response transformations

To quantify how optogenetic suppression altered auditory cortical responses, we compared neuronal firing rates during non-laser and laser conditions using population-level linear regression. Mean firing rates were calculated separately for non-laser and laser trials during clean target sound presentation by averaging responses across all stimulus repetitions and normalizing by stimulus duration to obtain mean firing rates (FR). Each neuron contributed one non-laser/laser firing rate pair for each of the four speaker locations, and observations were pooled across neurons to generate separate datasets for PV-Arch and SST-Arch animals.

Population response transformations were modeled using the linear relationship:

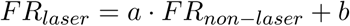

where the slope *a* reflects multiplicative (gain-like) changes in firing rate and the intercept *b* reflects additive (offset-like) changes. Model parameters were estimated using robust linear regression with a Random Sample Consensus (RANSAC) algorithm to reduce the influence of outlier observations exhibiting unusually large laser-induced firing-rate changes. Regression was performed using a first-order polynomial (degree = 1) model with a maximum of 1000 iterations and a confidence level of 99%. Observations were classified as inlier using a threshold equal to 5% of the range of firing rates in the laser condition, resulting in 124/160 and 108/136 inlier observations for the PV-Arch and SST-Arch datasets, respectively.

The fitted slope and intercept were used to characterize the dominant population response transformation. Slopes substantially greater than 1 with minimal intercept changes were interpreted as multiplicative (gain-like) transformations, whereas intercept shifts with slopes near unity were interpreted as additive (offset-like) transformations.

### Target-Masker Separation Analysis

For masked trials, target-to-masker speaker separation was quantified as the signed angular difference between the target and masker speaker locations:

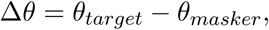

Speaker azimuths were referenced to the recorded right auditory cortex such that increasingly contralateral locations were assigned progressively more positive values. Consequently, positive target–masker separations correspond to spatial configurations in which the target was located at an equal or more contralateral position than the masker relative to the recorded hemisphere, whereas negative values indicate that the masker occupied the more contralateral location. This hemisphere-referenced convention follows the population-readout framework proposed by Maddox et al. (2012) and was used throughout analyses of masked neural discrimination.

### Statistics

All statistical analyses were performed in MATLAB using built-in statistical functions. Unless otherwise noted, statistical significance was assessed at *α* = 0.05, all t-tests were two-tailed, and data are reported as mean ± s.e.m.

For analysis comparing firing rate across speaker locations, firing rates were normalized within each neuron by dividing responses at each speaker location by the maximum firing rate observed during the non-laser condition. Paired *t*-tests were used to compare normalized firing rates between non-laser and optogenetic suppression conditions at each speaker location. Spatial tuning metrics, including modulation depth, equivalent rectangular receptive field (ERRF), response centroid, and sparseness were calculated for individual neurons under both non-laser and suppression conditions and compared using paired *t*-tests. Changes in firing rate for hotspot-containing neurons were analyzed using raw (non-normalized) firing rates averaged across speaker locations, allowing overall changes in neuronal responsiveness to be evaluated independently of spatial tuning. Unpaired *t*-tests were used only for comparisons between independent groups, including laminar analyses and quantification of Arch expression.

To evaluate neuronal discrimination across clean conditions and masked conditions, repeated-measures analysis of variance (RM-ANOVA) was performed with Condition (Non-laser versus Laser) and Spatial Configuration as within-neuron factors. For clean conditions, spatial configuration corresponded to the four target speaker locations (90°, 45°, 0°, −90°). For masked conditions, spatial configuration corresponded to positive target-to-masker separations (0°, 45°, 90°, 135°, 180°). Within each target-masker separation, discrimination values were averaged so that each neuron contributed a single observation per condition. Greenhouse-Geisser corrections were applied when assumptions of sphericity were violated, and Bonferroni-corrected post hoc comparisons were used for pairwise tests.

To determine whether optogenetic suppression altered the slope of the relationship between neuronal discrimination and target-masker separation (changing the slope), linear mixed-effects (LME) models were fitted using target-masker separation as a continuous predictor and experimental condition (Non-laser vs. Laser) as a fixed effect. Repeated-measurements from individual neurons were modeled using neuron-specific random intercepts and random slopes in MATLAB notation as:

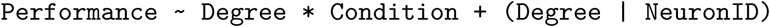

The primary statistical test was the interaction between target-masker separation and experimental condition with a significant interaction indicating that optogenetic suppression altered the relationship between spatial separation and neural discrimination. Throughout the manuscript, single, double, and triple asterisks denote *p* < 0.05, *p* < 0.01, and *p* < 0.001, respectively.

## Data availability

The data that support the findings of this study are available from the corresponding author upon reasonable request.

## Acknowledgements

We thank Rachael Bell for experiments support and Martín Irani-Cereceda for giving helpful comments on the manuscript. This research was supported by the National Science Foundation grants SMA-2319321 (H.J.G. and K.S.) and EFRI-2515342 (H.G.), as well as a Brain Research Foundation Award BRFSG-2023-13 (H.G.).

## Author contributions

Z.Q. and J.C.N. performed all experiments. Z.Q. and J.C.N. analyzed the data. H.J.G. and K.S. obtained funding and supervised the study. Z.Q., H.J.G., and K.S. wrote the manuscript and contributed to the interpretation of the results.

## Supplementary Information

**Supplementary Figure 1.**
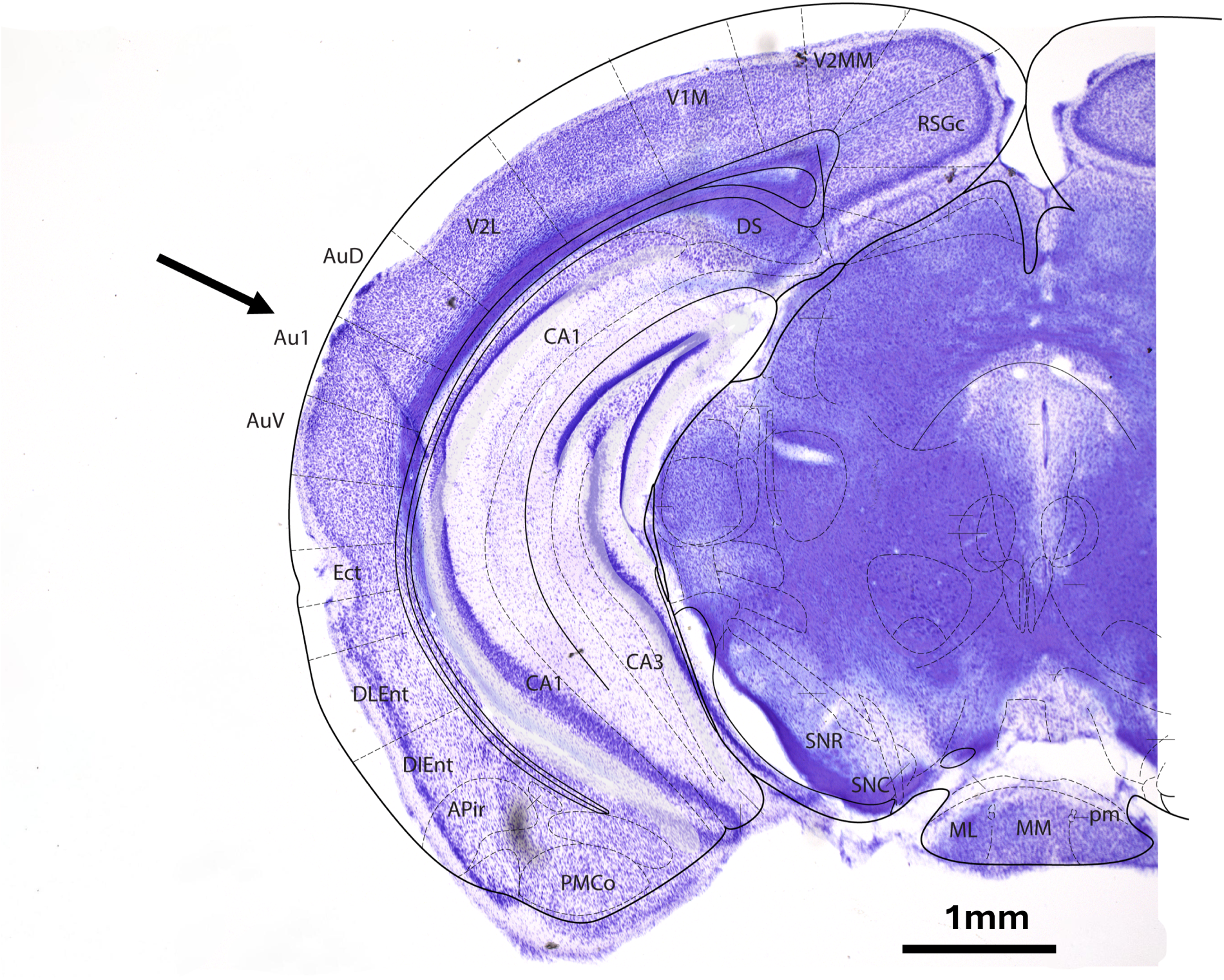
Histological verification of electrode placement in primary auditory cortex. Representative Nissl-stained coronal section showing the recording electrode track within the right primary auditory cortex (A1). The arrow indicates the location of electrode placement following completion of the experiments.

**Supplementary Figure 2.**
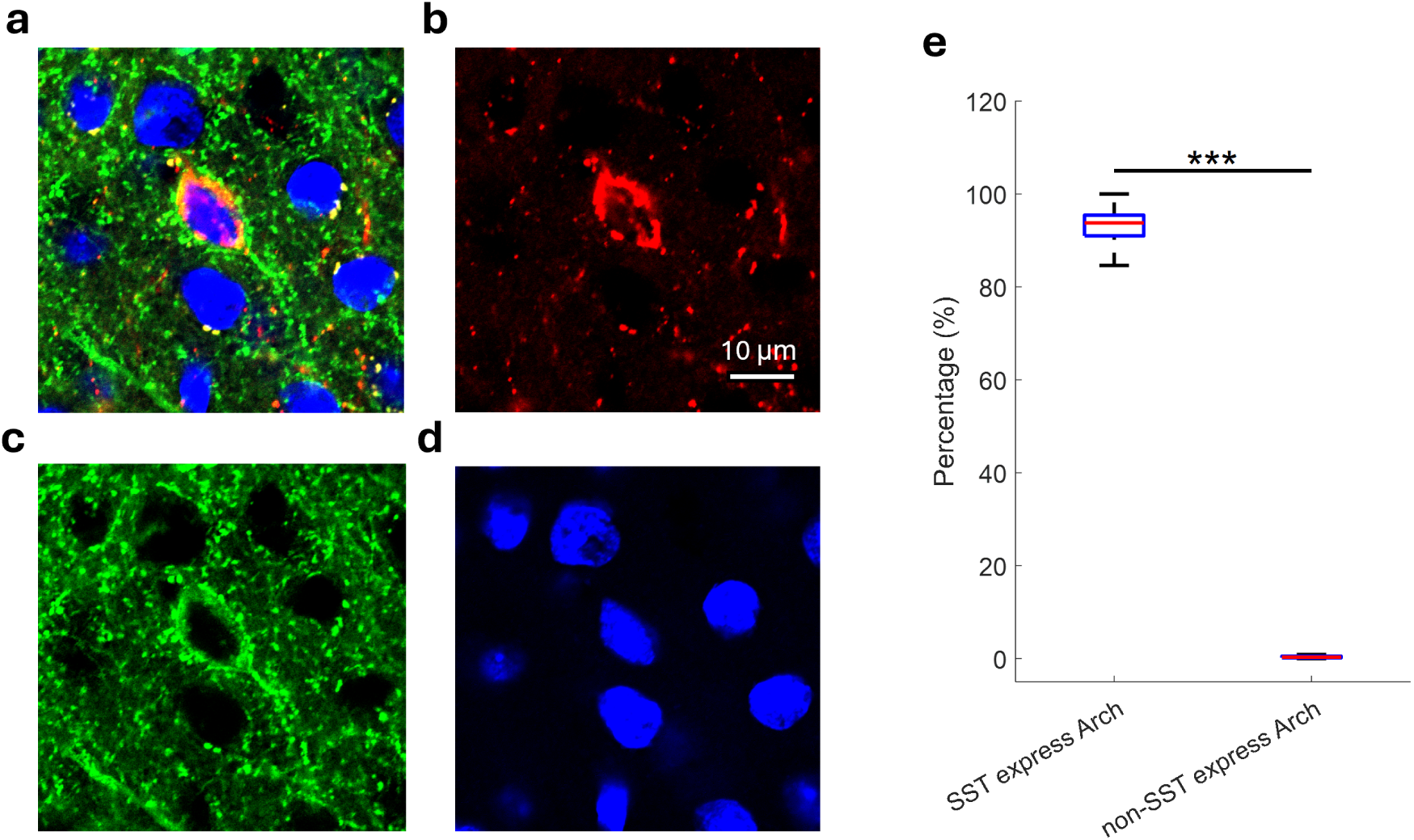
Histological validation of SST-targeted Arch expression. Arch expression in PV-Arch mice was validated previously (Nocon et al. 2023a); therefore, only SST-Arch animals are shown here. **a**. Representative fluorescence image from an SST-Arch mouse illustrating Arch-GFP expression within the auditory cortex. **b–d**. Corresponding single-channel images showing SST immunolabeling (b, red), Arch–GFP expression (c, green), and DAPI nuclear labeling (d, cyan). **e.** Quantification of Arch expression specificity in SST-Arch animals (n = 8 mice). The majority of Arch-GFP-positive neurons colocalized with SST immunoreactivity, confirming selective expression of SST interneurons (unpaired t-test, p = 4.5997× 10^-18^; see Methods).

**Supplementary Figure 3.**
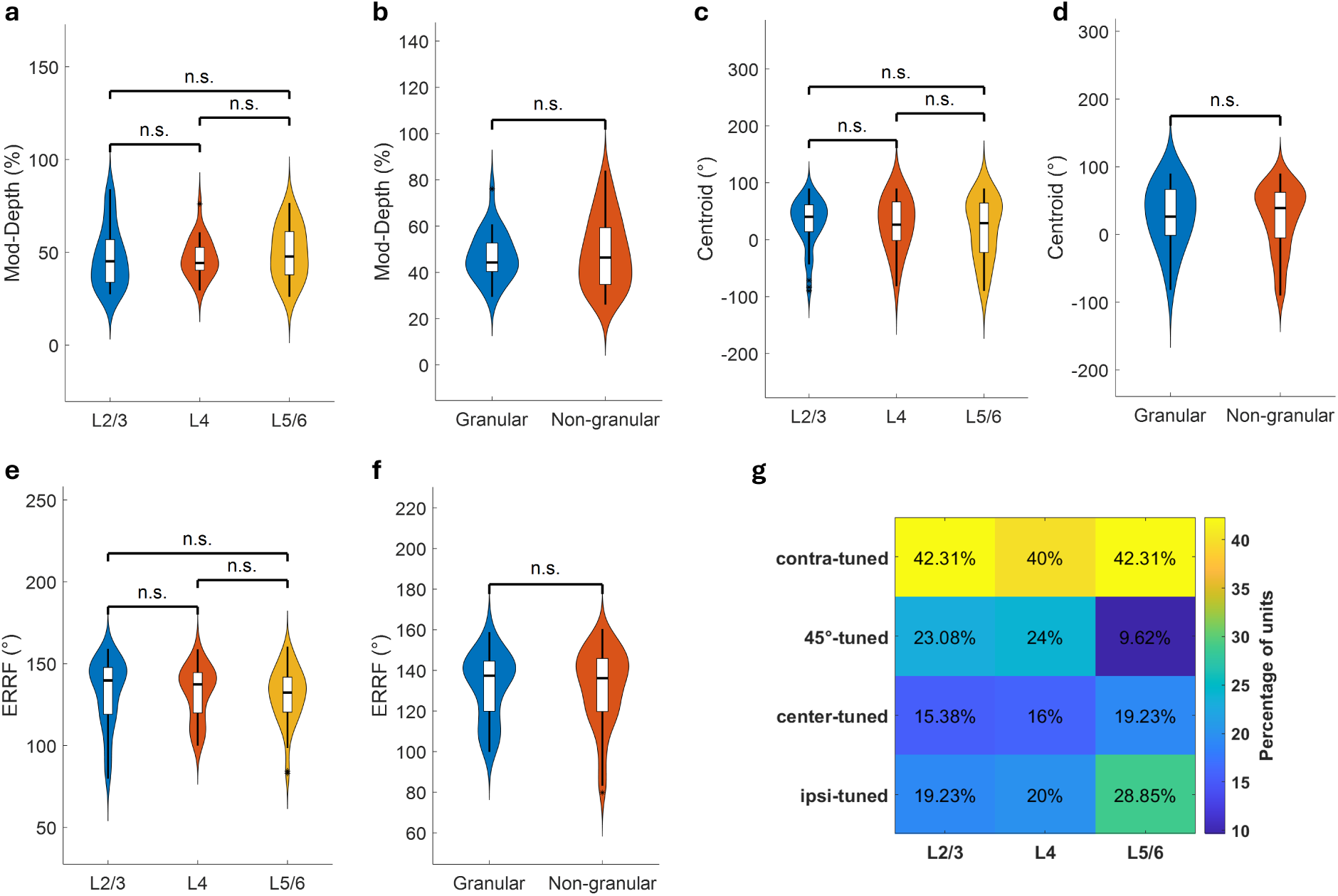
Spatial tuning properties are conserved across cortical layers. Spatial tuning metrics and the distribution of spatial tuning types are shown across cortical layers. A total of 137 auditory spatial units were recorded. For 8 units, laminar assignment could not be determined because the initial current sink in the CSD profile was too small to be reliably identified. The remaining 129 units were assigned to cortical layers based on CSD analysis. **a, c, e**. Modulation depth (a), response centroid (c), and equivalent rectangular receptive field (ERRF) width (e) for neurons assigned to layers 2/3 (n = 52), 4 (n = 25), and 5/6 (n = 52). **b, d, f**. Corresponding comparisons between granular (L4; n=25) and non-granular layers (n=104), where L2/3 and L5/6 were grouped as non-granular. All comparisons used unpaired t-tests and revealed no significant differences (all p > 0.05). **g**. Distribution of preferred spatial tuning classes across cortical layers. No significant differences in modulation depth, centroid position, or ERRF were observed across cortical layers, indicating that the heterogeneous but contralaterally biased representation of auditory space described in Figure 2 is broadly preserved throughout A1.

**Supplementary Figure 4.**
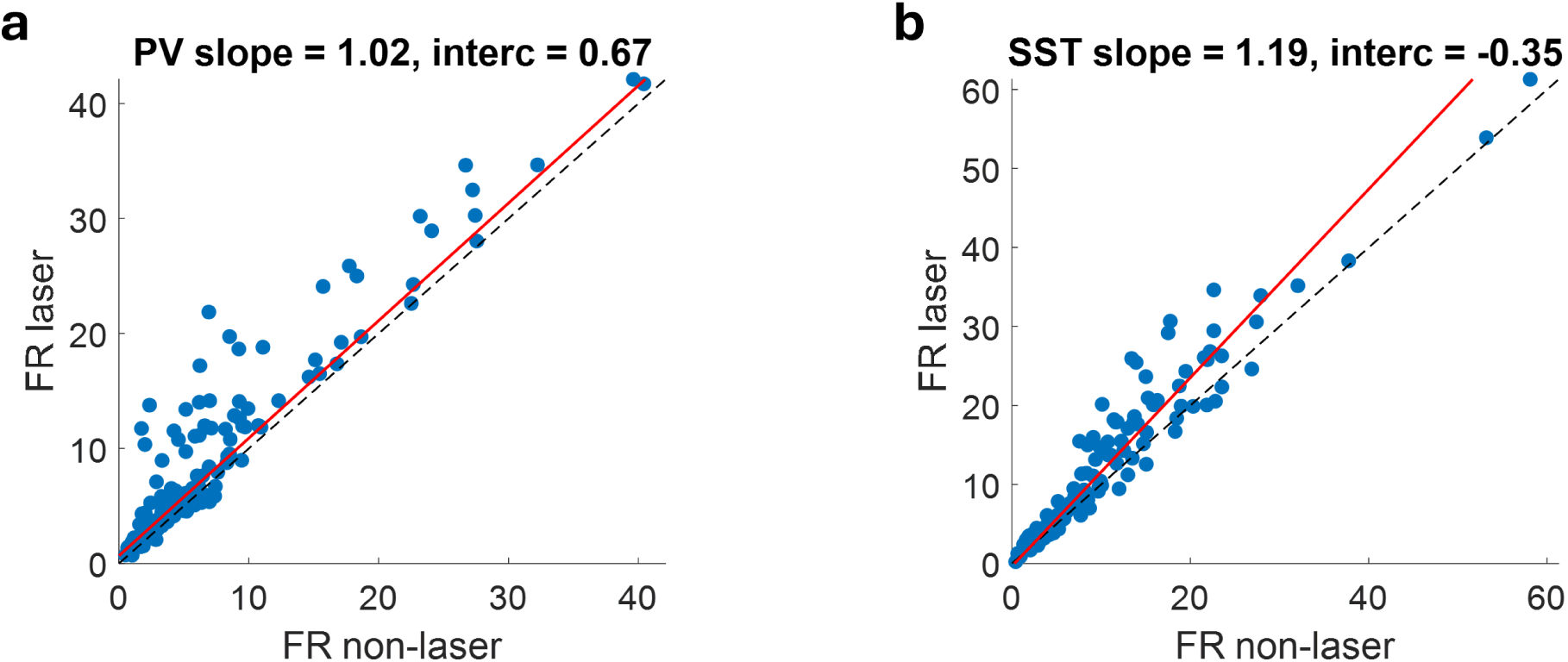
PV and SST suppression produce distinct population input-output transformations. **a.** Population relationship between unit firing rates (FR) measured during non-laser and PV suppression across all unit–location observations recorded from PV-Arch animals. Each point represents the mean firing rate for one neuron at one speaker location during clean target presentation. The red line shows the robust linear regression (RANSAC) fit (slope = 1.02, intercept = 0.67), indicating that PV suppression primarily shifted the input–output relationship through an additive/subtractive (offset-like) transformation (n = 160). **b**. Corresponding analysis for SST-Arch animals. The fitted regression (slope = 1.19, intercept = −0.35) indicates that SST suppression predominantly altered the slope of the input–output relationship, consistent with a divisive/multiplicative (gain-like) transformation (n = 136). These analyses quantify the distinct population response transformations underlying the differences in spatial tuning. Here, *n* denotes the number of unit–location observations: each unit contributes four FR values measured at four speaker locations during clean target presentation.

**Supplementary Figure 5.**
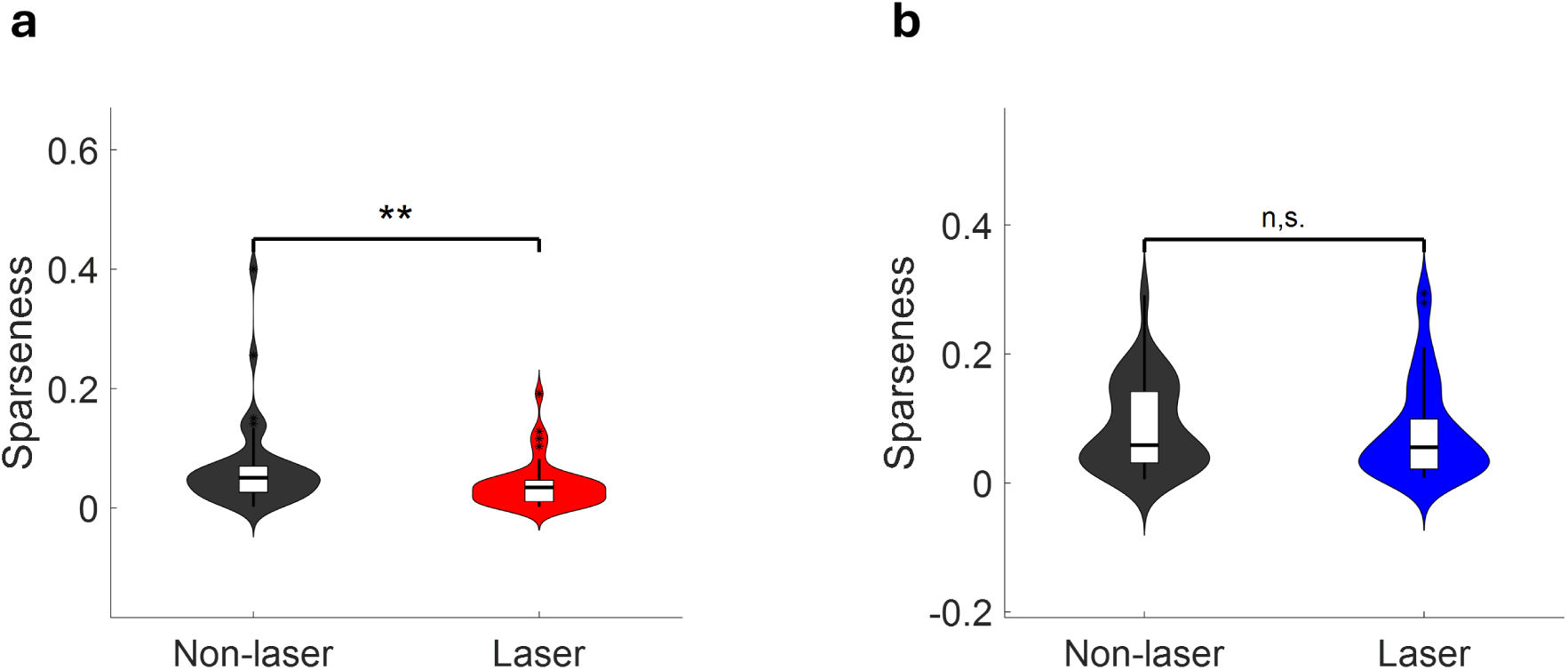
PV and SST suppression differentially alter the sparseness of auditory responses. **a**. Sparseness of auditory cortical responses across the four speaker locations during clean target presentation in PV-Arch animals (n = 40). Each point represents one spatially tuned neuron measured during non-laser and PV suppression. PV suppression significantly reduced response sparseness, indicating broader responses across speaker locations (paired t-test, p = 0.001). **b**. Corresponding analysis for SST-Arch animals (n=34). SST suppression did not significantly alter the firing-rate sparseness across four speaker locations indicating that response breadth remained largely unchanged despite optogenetic suppression.

**Supplementary Figure 6.**
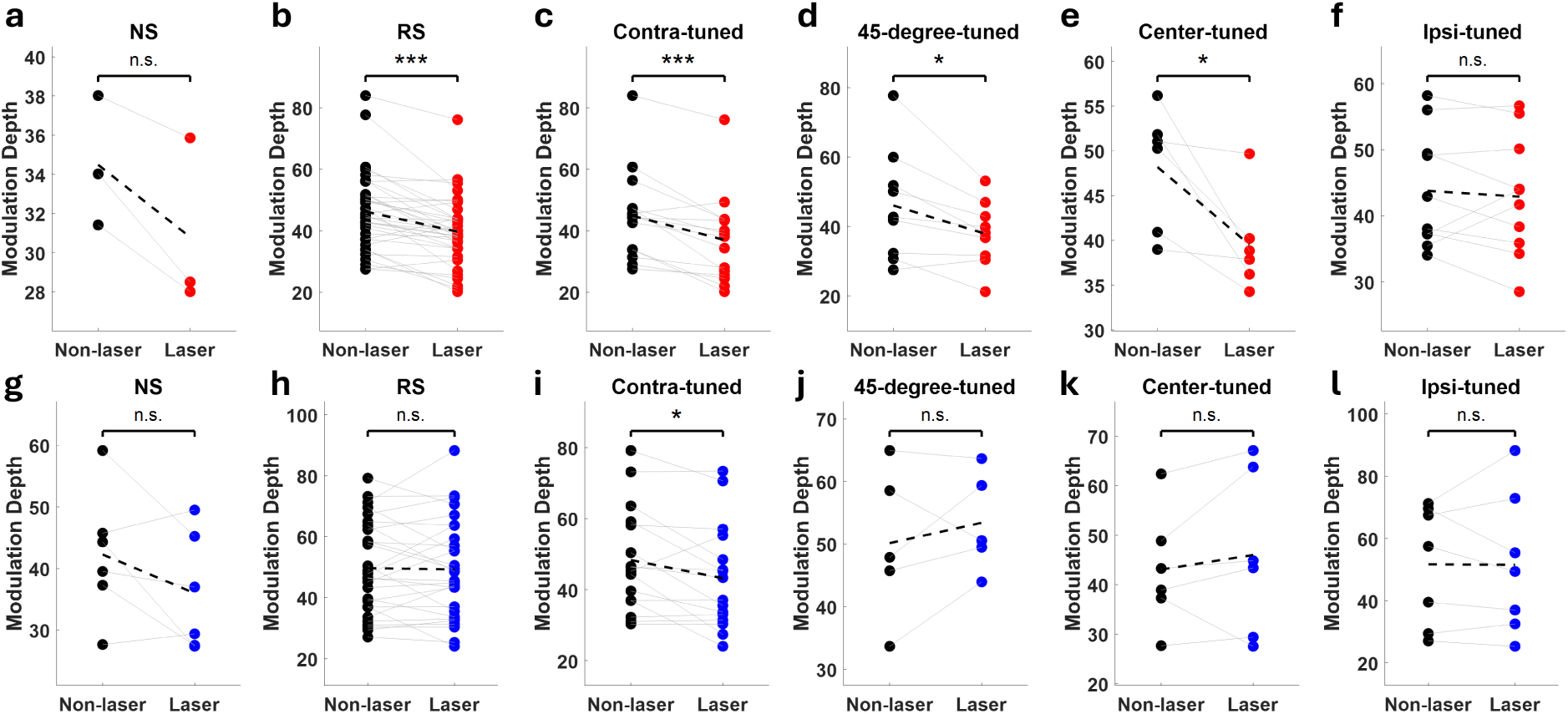
PV and SST suppression differentially affect modulation depth across neuronal subpopulations during optogenetic silencing. Changes in modulation depth across neuronal subpopulations classified either by spike waveform or by preferred spatial tuning. For waveform-based classification, units with a peak-to-trough duration < 0.5 ms were classified as narrow-spiking (NS), whereas units with longer durations were classified as regular-spiking (RS). **a–f.** PV-Arch animals (red) modulation depth measured during non-laser and PV-suppression trials for (a) NS units (n = 3), (b) RS units (n = 37, p = 4.03 × 10^-6^), (c) contra-tuned units (n = 15, p = 3.53 × 10^-4^), (d) 45°-tuned units (n = 9, p = 0.017), (e) center-tuned units (n = 6, p = 0.027), and (f) ipsi-tuned units (n = 10). **g–l.** Corresponding analyses for SST-Arch animals (blue). Across neuronal subpopulations, PV suppression consistently reduced modulation depth, whereas SST suppression of (g) NS units (n = 6) and (h) RS units (n = 28) produced comparatively small and less consistent effects, with only (i) contra-tuned units showing a significant change associated with optogenetic suppression (n = 16, p = 0.019); (j) 45°-tuned units (n = 5), (k) center-tuned units (n = 6), and (l) ipsi-tuned units (n = 7) showed no significant change.

**Supplementary Figure 7.**
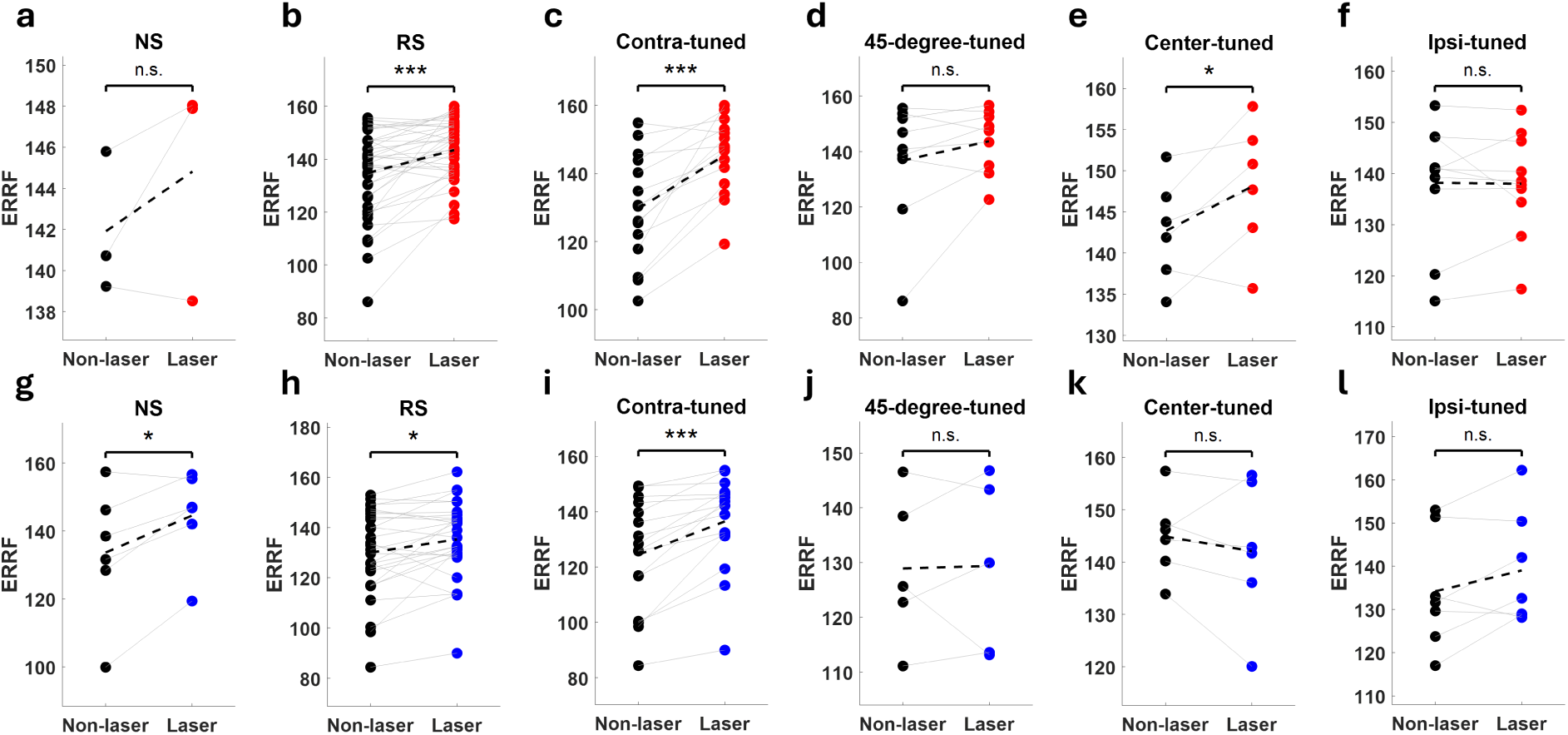
PV and SST suppression differentially affect the equivalent rectangular receptive field (ERRF) across neuronal subpopulations during optogenetic silencing. Equivalent rectangular receptive field (ERRF) width, a measure of spatial tuning breadth, was quantified across neuronal subpopulations classified by spike waveform or preferred spatial tuning. **a–f.** PV-Arch animals (red). ERRF measured during non-laser and PV suppression for (a) narrow-spiking (n=3), (b) regular-spiking n = 37, p = 7.74 × 10^-5^), (c) contralateral-tuned (n = 15, p = 8.45 × 10^-5^), (d) 45°-tuned (n = 9), (e) center-tuned (n = 6, p = 0.047), and (f) ipsilateral-tuned neurons (n = 10). **g–l**. Corresponding analyses for SST-Arch animals (blue). Across neuronal subpopulations, PV suppression increased ERRF width, whereas SST suppression of (g) NS (n = 6, p = 0.019) and (h) RS units (n = 28, p = 0.013) produced a more selective change in tuning breadth with only (i) contra-tuned units showing a significant change associated with optogenetic suppression (n = 16, p = 1.10 × 10^-4^); (j) 45°-tuned units (n = 5), (k) center-tuned units (n = 6), and (l) ipsi-tuned units (n = 7) showed no significant change.

**Supplementary Figure 8.**
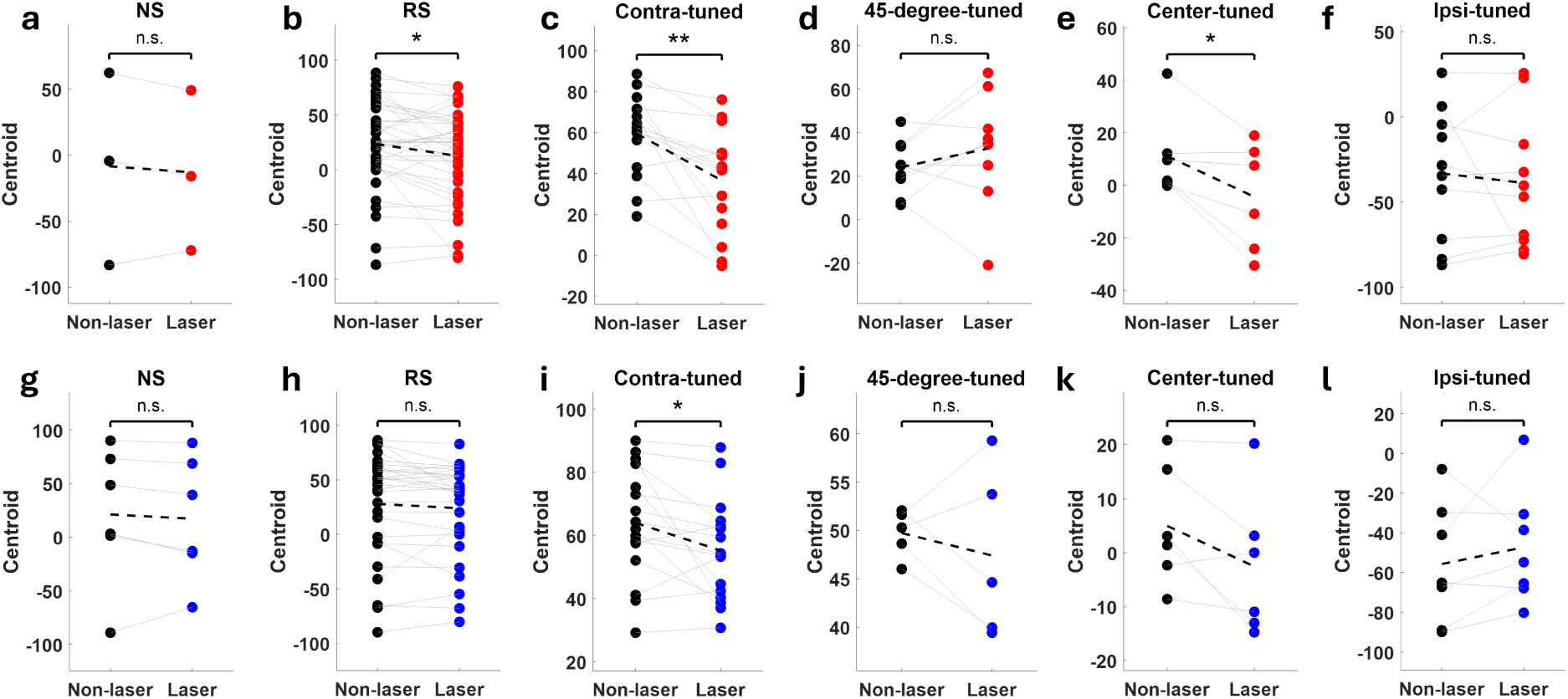
PV and SST suppression differentially affect response centroid across neuronal subpopulations. Response centroid, a measure of preferred sound location, was quantified across neuronal subpopulations classified by spike waveform or preferred spatial tuning. **a–f.** Response centroid measured during non-laser and PV suppression for (a) narrow-spiking (n = 3), (b) regular-spiking (n = 37, p = 0.019), (c) contralateral-tuned (n = 15, p = 0.001), (d) 45°-tuned (n = 9), (e) center-tuned (n = 6, p = 0.035), and (f) ipsilateral-tuned neurons (n = 10) recorded from PV-Arch animals (red). **g–l.** Corresponding analyses measured during non-laser and SST suppression for (g) narrow-spiking (n = 6), (h) regular-spiking (n = 28), (i) contralateral-tuned (n = 16, p = 0.031), (j) 45°-tuned (n = 5), (k) center-tuned (n = 6), and (l) ipsilateral-tuned neurons (n = 7) in SST-Arch animals (blue). PV suppression produced broader changes in response centroid across neuronal subpopulations, whereas SST suppression resulted in comparatively smaller shifts in preferred sound location.

**Supplementary Figure 9.**
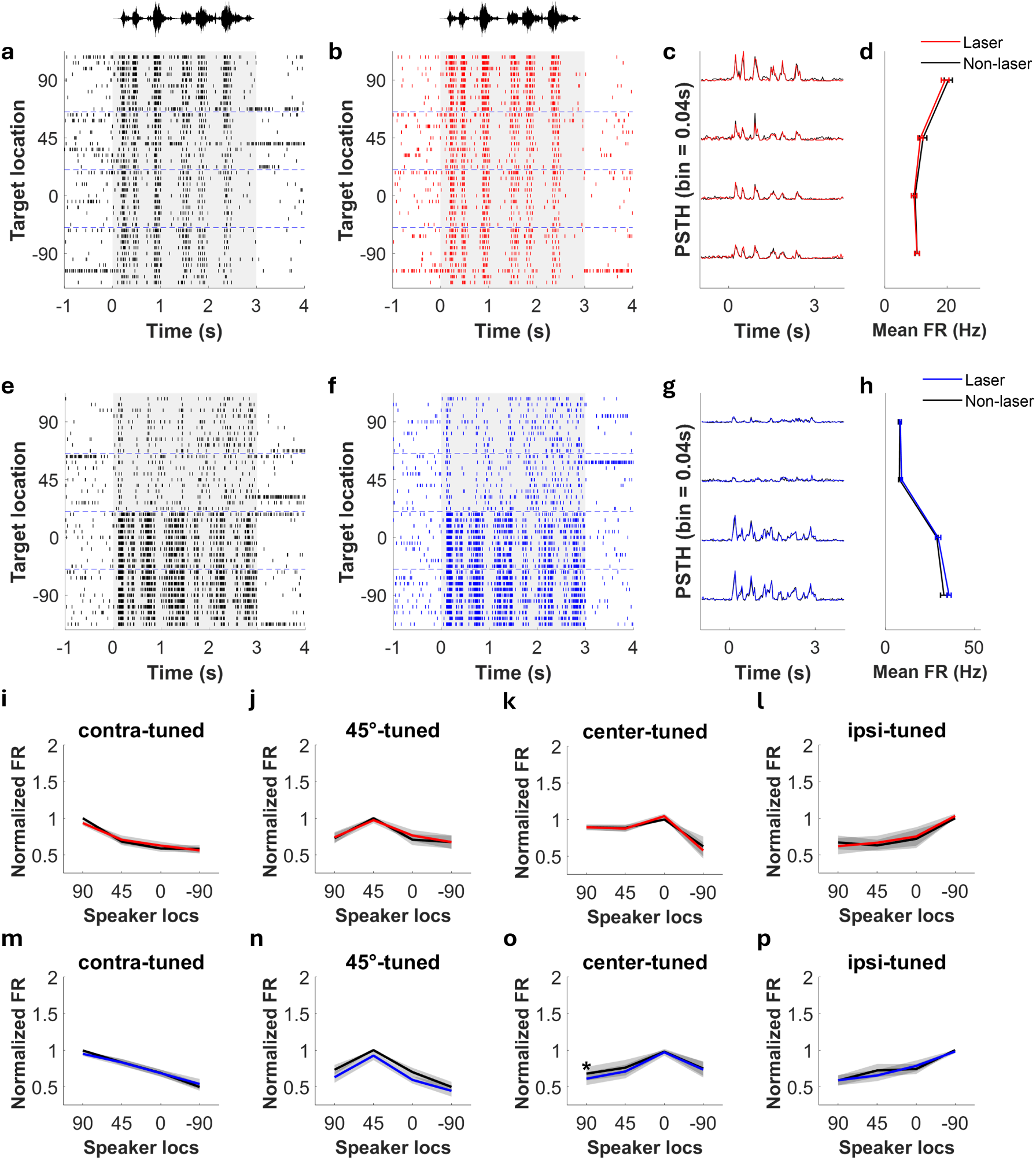
Laser illumination alone does not alter auditory spatial tuning. **a–d**. Representative auditory cortical neuron recorded from a PV-Cre control mouse during non-laser (a: black) and laser (b: red) trials. Peri-stimulus time histogram (c) and spatial tuning curves (d) show minimal changes during laser illumination compared to non-laser trials. **e–h**. Corresponding example neuron recorded from an SST-Cre control mouse. **i–l**. Population-averaged spatial tuning curves grouped according to preferred sound location (contralateral-tuned, n = 12; 45°-tuned, n = 6; center-tuned, n = 4; ipsi-tuned, n = 5) for PV-Cre control mice. **m–p**. Corresponding population tuning curves for SST-Cre control mice grouped by tuning type (contralateral-tuned, n = 11; 45°-tuned, n = 6; center-tuned, n = 6; ipsi-tuned, n = 8). Across both control genotypes, laser illumination alone produced no systematic changes in auditory spatial tuning, indicating that the alterations observed in Arch-expressing animals require optogenetic suppression of inhibitory interneurons rather than nonspecific optical stimulation. Note the small exception for center-tuned units which differed at the 90° speaker location (p = 0.016).

**Supplementary Figure 10.**
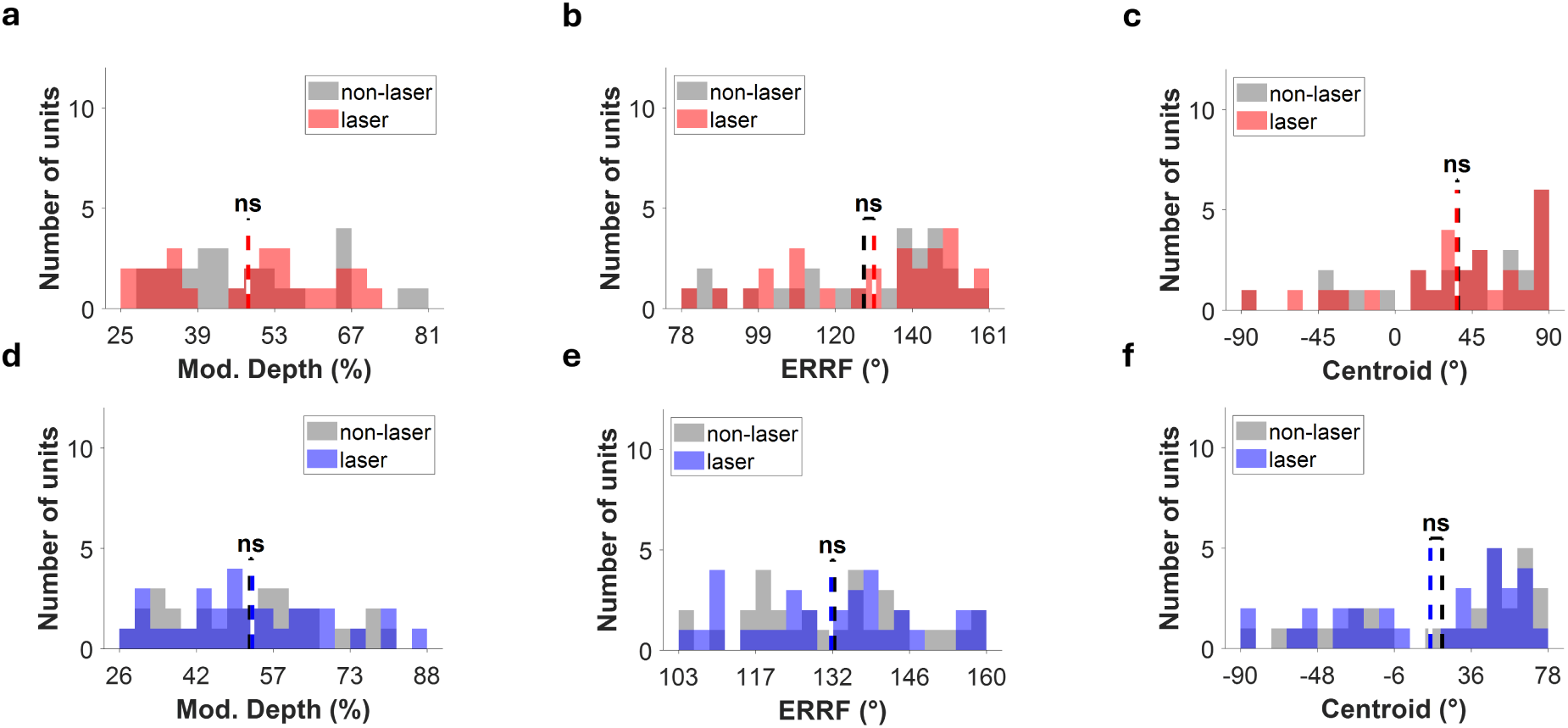
Laser illumination alone does not alter spatial tuning metrics. To assess the effect of laser illumination independent of optogenetic suppression, modulation depth, equivalent rectangular receptive field (ERRF), and response centroid were compared between non-laser and laser trials in PV-Cre and SST-Cre mice lacking Arch expression. **a–c**. Population distributions of modulation depth (a), ERRF width (b), and response centroid (c) for spatially tuned neurons recorded from PV-Cre control mice (n = 27). No significant differences were observed between non-laser(gray) and laser (red) conditions. **d–f**. Corresponding analysis for spatially tuned neurons from SST-Cre control mice (n = 31) in non-laser (gray) and laser (blue) sessions.

**Supplementary Figure 11.**
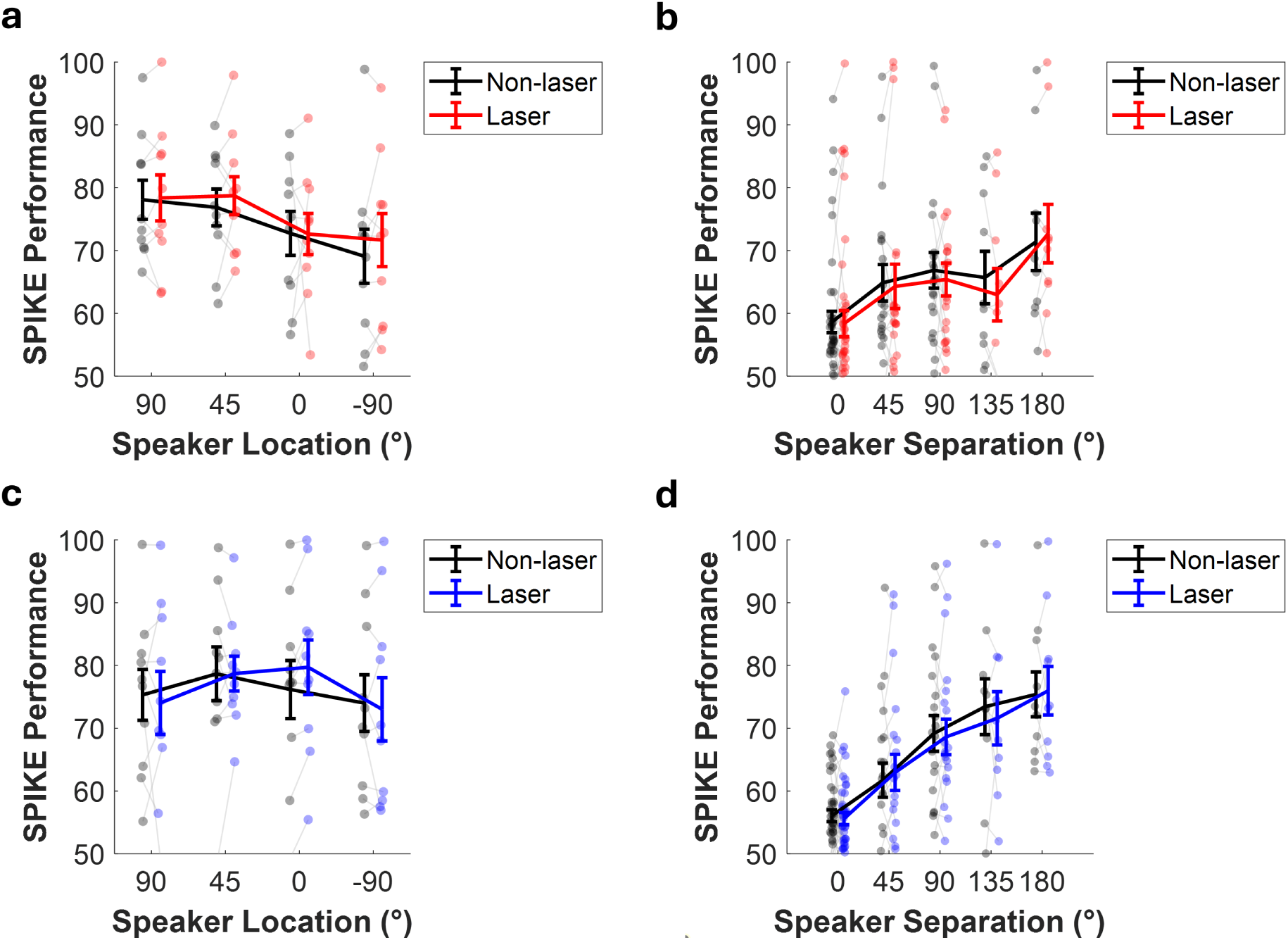
Laser illumination alone does not alter neural discriminability in control animals. Effects of laser illumination alone on SPIKE-distance-based neural discrimination in PV-Cre and SST-Cre control mice lacking Arch expression. **a-b**. Neural discrimination during clean (a) and masked (b) listening conditions for hotspot-containing neurons recorded from PV-Cre control animals. Laser illumination did not significantly alter discrimination under either condition. **c-d.** Corresponding analyses for SST-Cre control animals. Similarly, laser illumination produced no significant changes in clean (c) or masked (d) neural discrimination. In masked listening conditions, laser illumination also failed to alter the relationship between target-to-masker separation and neural discrimination (linear mixed-effects model, interaction).

**Supplementary Figure 12.**
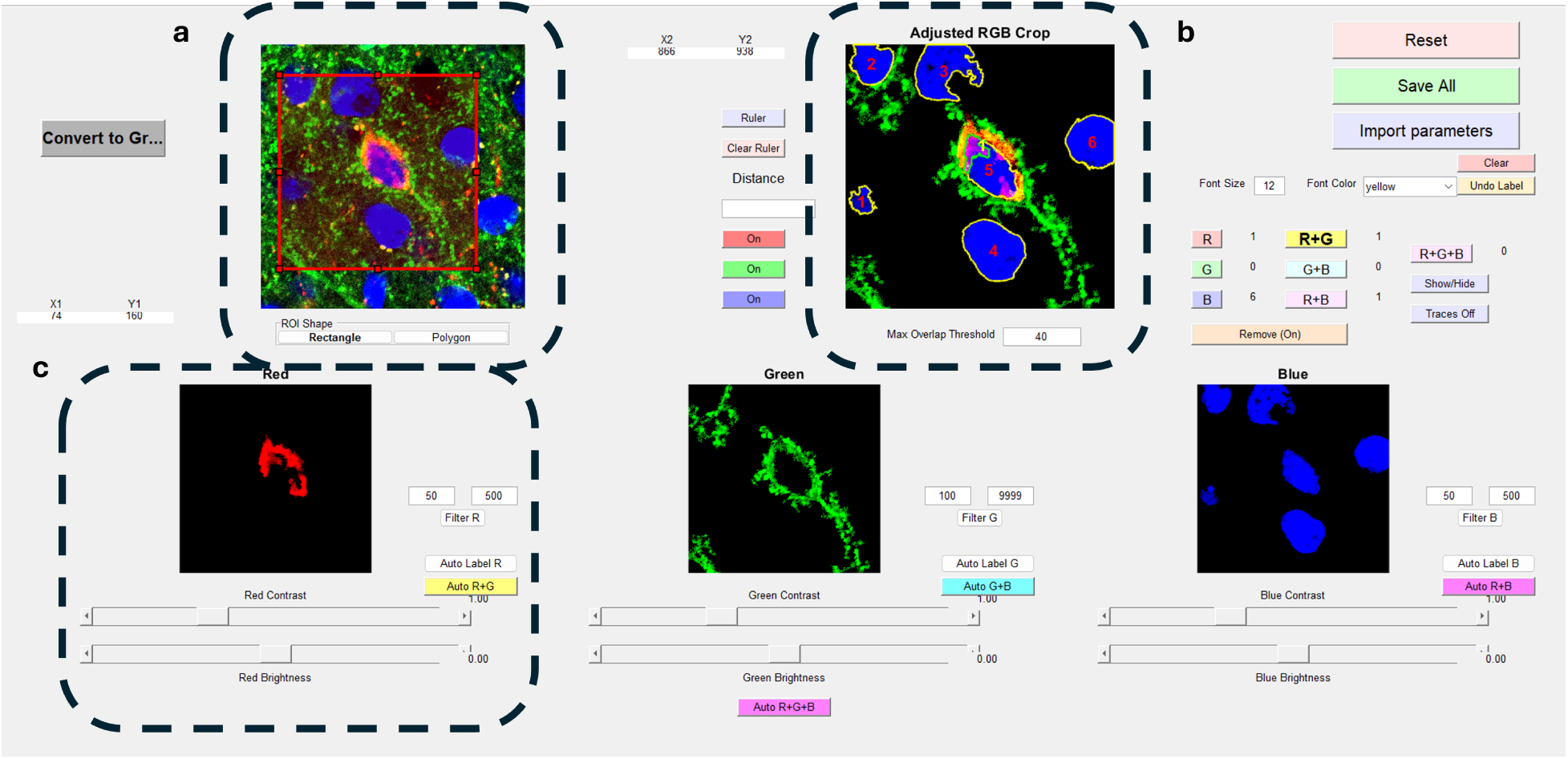
SliceQuant workflow for fluorescence image quantification. Use of SliceQuant MATLAB-based graphical user interface developed to standardize quantification of fluorescence images used for assessing Arch expression and cell-type specificity (see Methods). **a.** Interactive region-of-interest (ROI) selection. Users define rectangular or polygonal ROIs for subsequent image analysis. **b.** Example analysis output following ROI selection, image filtering, and automated cell detection. Detected neurons can be manually reviewed and edited before exporting results. **c.** Channel-specific image-processing interface showing adjustable brightness and contrast, area-based filtering, and automated cell-labeling parameters used during fluorescence quantification.

